# DNA clamp function of the mono-ubiquitinated Fanconi Anemia FANCI-FANCD2 complex

**DOI:** 10.1101/854133

**Authors:** Renjing Wang, Shengliu Wang, Ankita Dhar, Christopher Peralta, Nikola P. Pavletich

## Abstract

The FANCI-FANCD2 (ID) complex, mutated in the Fanconi Anemia (FA) cancer predisposition syndrome, is required for the repair of replication forks stalled at DNA interstrand crosslinks (ICL) and related lesions^1^. The FA pathway is activated when two replication forks converge onto an ICL^2^, triggering the mono-ubiquitination of the ID complex. ID mono-ubiquitination is essential for ICL repair by excision, translesion synthesis and homologous recombination, but its function was hitherto unknown^1,3^. Here, the 3.48 Å cryo-EM structure of mono-ubiquitinated ID (ID^Ub^) bound to DNA reveals that it forms a closed ring that encircles the DNA. Compared to the cryo-EM structure of the non-ubiquitinated ID complex bound to ICL DNA, described here as well, mono-ubiquitination triggers a complete re-arrangement of the open, trough-like ID structure through the ubiquitin of one protomer binding to the other protomer in a reciprocal fashion. The structures, in conjunction with biochemical data, indicate the mono-ubiquitinated ID complex looses its preference for ICL and related branched DNA structures, becoming a sliding DNA clamp that can coordinate the subsequent repair reactions. Our findings also reveal how mono-ubiquitination in general can induce an alternate structure with a new function.

---

FANCI and FANCD2 are paralogs that bind to DNA with preference for branched structures including Holliday junction, overhang and replication fork DNA^4-7^. The previous crystal structure of the mouse ID complex showed that it forms an open trough-like structure with two basic grooves, one on each paralog^7^. A 7.8 Å crystallographic map of FANCI bound to splayed Y DNA confirmed that its basic groove is the site of dsDNA binding, and also identified a likely single-stranded DNA binding region^7^. However, it has not been clear how these DNA binding activities relate to the function of the ID complex in replication and ICL repair. The mouse ID structure also showed the mono-ubiquitination sites are embedded inside the FANCI-FANCD2 interface^7^, but did not shed light on the function of mono-ubiquitination.

To address these questions, we first collected cryo-EM data on the human, full-length ID complex bound to an ICL-containing DNA constructed by crosslinking two modified oligonucleotides with a triazole moiety^8^ (Fig. 1a). This ICL DNA mimics two replication forks converging on an ICL, an event shown to activate the FA pathway^2^. The initial consensus reconstruction with 231,943 particles extended to 3.40 Å, as determined by the gold-standard fourier shell correlation (FSC) procedure^9^ (Extended Data Figs 1a and b). The map showed that the human ID complex has an overall structure and FANCI-FANCD2 interface very similar to the mouse ID complex^7^, with each paralog consisting of N-terminal helical repeats that form an extended α−α solenoid (henceforth NTD, residues 1 to 550 of FANCI and 45 to 587 of FANCD2), followed by a helical domain (HD) that reverses the direction of the solenoid, and a C-terminal helical repeat domain (CTD, residues 805 to 1280 of FANCI and 929 to 1376 of FANCD2) (Figs 1a and b). The consensus reconstruction showed clear density for FANCI and its bound dsDNA, and density for single-stranded DNA extending from the FANCI-bound duplex, both approximately where they were seen in the mouse FANCI-Y DNA crystallographic map^7^. FANCD2 had clear density for its NTD, but poor density for its CTD and what appeared to be a DNA duplex bound to it (Extended Data Fig. 1b). 3D classification indicated that the FANCD2 CTD exhibited substantial conformational flexibility, its relative position moving by up to 23 Å due to rotation within the HD domain (starting around residue 645) (Extended Data Figs 1c and d). FANCI did not exhibit this flexibility, as its NTD contains a helical protrusion that packs with and stabilizes the CTD and which is absent from FANCD2^7^. To better account for the flexibility of FANCD2, we used multi-body refinement with RELION-3^10^. This improved the solvent-corrected resolution of FANCD2 CTD and its associated dsDNA to 3.77 Å, and the remainder of the complex to 3.32 Å (Extended Data Fig. 1a; density shown in Extended Data Figs 2a to c).

**Figure 1.**
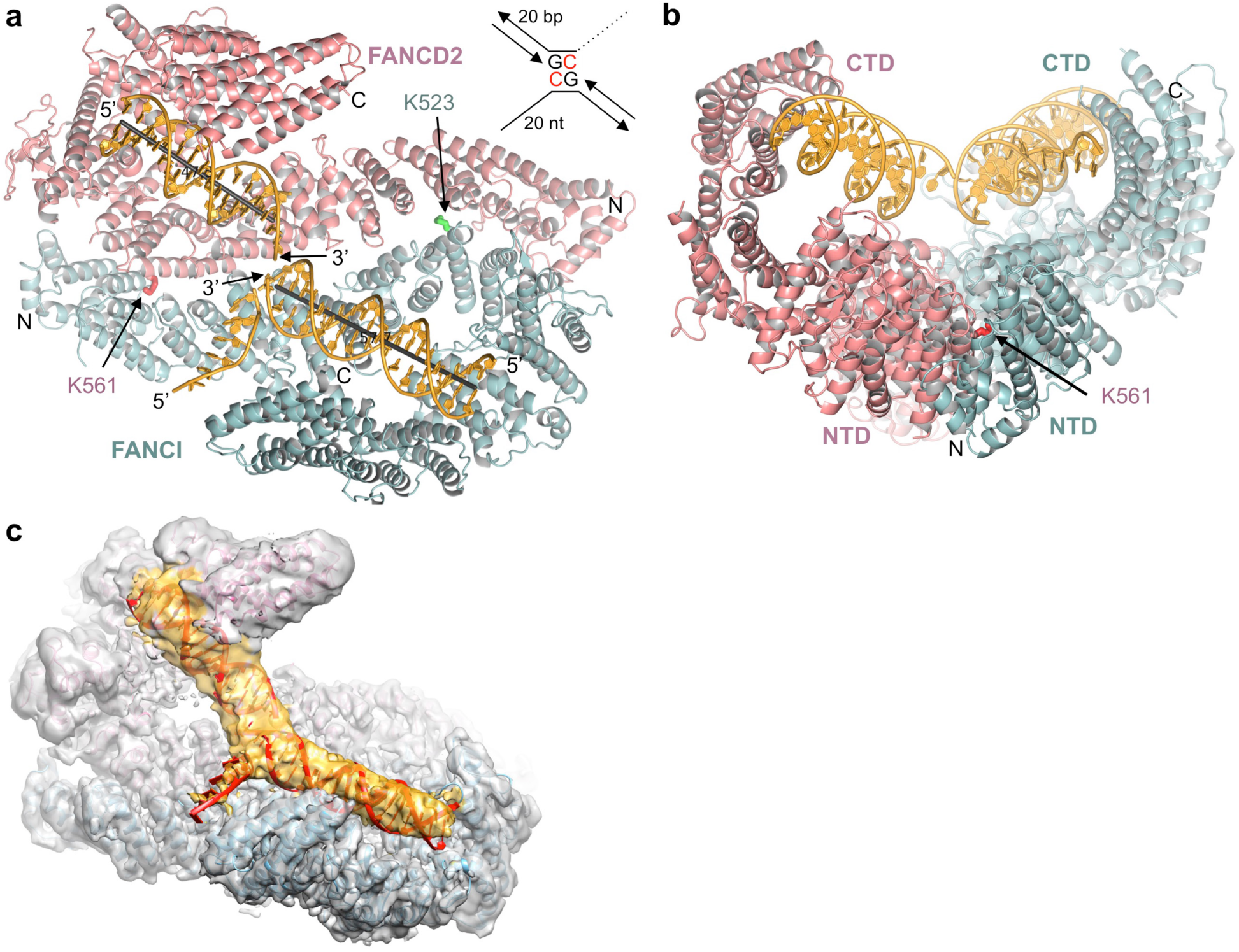
Overall Structure of the human ID complex bound to ICL DNA. **a**, Cartoon representation with FANCI colored cyan, FANCD2 salmon, and ICL DNA yellow. The mono-ubiquitination site Lys523 and Lys561 are shown as sticks (red for FANCD2, green for FANCI). The 5’ and 3’ ends of the two DNA duplexes and of the FANCI-bound ssDNA are labeled. The helical axes of the duplexes are indicated as gray lines. The N and C termini are labeled. Inset shows schematic of the ICL DNA, with the deoxycytidine residues crosslinked by a triazole in red (not shown in structure). **b**, View looking down the left side of the horizontal axis of **a. c**, Cryo-EM density prior to post-processing. Orientation similar to **a**. DNA is red, and its density yellow.

The improved maps showed continuous double-stranded DNA density extending from the FANCI groove to the FANCD2 groove (Fig. 1c). The DNA is sharply kinked near the center, where the ICL would be based on the lengths of the flanking duplexes. There was no density that could correspond to the 5’ overhang ssDNA of the FANCD2-bound duplex. We modeled the overall DNA with an 18 base pair (bp) duplex and an 8 nucleotide ssDNA bound to FANCI, and a 15 bp duplex on FANCD2, and refined the model with the composite map option of REFMAC5^11^ (Extended Data Table 1). We have not modeled the ICL and its immediate surroundings that include the link between the FANCI duplex and its 5’ overhang ssDNA due to the overall high temperature factors and limited resolution of the DNA density. In the refined model, the FANCI and FANCD2-bound duplexes are at a ∼33° angle. Their helical axes are non-collinear, being dislocated laterally by ∼14 Å (Fig. 1a). This non-collinear arrangement of the two duplexes is largely unaffected by the conformational flexibility of the FANCD2 CTD, as determined by principal component analysis of the multi-body angles and refinement of the model against reconstructions representing the range of motion (Extended Data Fig. 3, and Extended Videos 1 and 2).

FANCI binds to DNA using an extended basic groove consisting of parts of the NTD, HD and CTD domains. A 4 bp portion of dsDNA distal from the ICL is bound by a semi-circular constriction between the HD and CTD domains, while the rest of the duplex is bound by the NTD, and the ssDNA runs across the last two helical repeats of the CTD (Extended Data Figs 2c and 4a). FANCD2’s DNA binding activity diverged from that of FANCI, as the HD portion of its semi-circular groove is acidic and is uninvolved in DNA binding, and the FANCD2 groove is truncated owing to the absence of the NTD-CTD link that extends the base of the groove in FANCI (Extended Data Figs 4b to d). Rather, the ICL-distal portion of the DNA duplex is bound by a localized basic patch on the FANCD2 CTD, largely non-overlapping with the DNA-binding surfaces of FANCI. Here, an arginine side chain inserts into the minor groove of the DNA, flanked by contacts to the backbone of the two DNA strands (Arg1352, Extended Data Figs 4b and c). The arginine minor groove insertion is associated with compression of the minor groove, a common arrangement in DNA-binding proteins.

The ID complex can associate with replication forks independent of ICLs^12-14^, and it is implicated in replication fork recovery after stalling caused by genomic stress or unstable genomic loci^3,15^. We thus also evaluated ID binding to DNA structures that can arise during replication. We collected cryo-EM data with 5’ flap, reversed fork-like Holliday junction (HJ), and replication fork DNA. While a FANCI-bound duplex was evident in all three maps, there was essentially no density for a FANCD2-bound duplex either in the consensus reconstructions or in individual 3D classifications or their subsequent auto-refinement (Extended Data Figs 5 to 7). This suggests that FANCD2 engagement requires a translocation of the helical axes of the two duplexes, because while the 5’ flap, HJ and fork DNA structures have a discontinuity in the DNA backbone, they remain stacked^16,17^, and translocating their helical axes would impose an energetic penalty.

Together, these data indicate that the principal dsDNA binding activity resides with FANCI, with canonical dsDNA sufficing for FANCI binding. This may account for observations that FANCI alone, but not FANCD2 alone accumulates at active replication forks before stalling^14^. It is also consistent with the ID complex exhibiting only modest specificity for branched DNA structures in biochemical assays^5,7^ (dissociation constants of ∼12, ∼10, ∼20 and ∼24 nM, respectively, for ICL, HJ, fork and 5’ flap DNA compared to ∼45 nM for dsDNA; Extended Data Fig. 7f). We presume that once FANCD2 engages in DNA binding at an ICL or a related DNA structure, this would likely prevent the ID complex from laterally diffusing along the DNA, stabilizing it at the lesion.

We next sought to investigate the function of mono-ubiquitination by determining the structure of the mono-ubiquitinated ID. For this, we constructed a stably-transfected HEK-293F cell line overexpressing eight subunits of the FA Core complex ubiquitin ligase (FANC A, B, C, E, F, G, L, FAAP100) (Extended Data Fig. 8a). Using the purified FA core complex with the FANCT/UBE2T E2 and E1, we assayed for the ubiquitination of the ID complex in the presence of ICL DNA or a variety of other DNA molecules previously shown to promote FANCD2 ubiquitination in a FANCI-dependent manner^18-20^ (Extended Data Fig. 8b and c). We purified the reaction products on size exclusion chromatography and found that the mono-ubiquitinated ID complex (henceforth ID^Ub^) remained bound to ICL DNA, in contrast to the non-ubiquitinated ID complex that retained very little DNA under the same conditions (Extended Data Figs 8d and e). We collected cryo-EM data of ID^Ub^ bound to four different DNA molecules: ICL DNA, 5’ flap DNA, nicked DNA and canonical dsDNA. The largest data set was collected with nicked DNA, as this gave the highest yield in the mono-ubiquitination reaction. The 3D auto-refinement of 301,058 particles led to a reconstruction extending to 3.57 Å resolution, as determined by the gold-standard fourier shell correlation (FSC) procedure (Extended Data Figs 9a and b). We calculated focused reconstructions with three partially overlapping masks (3.44, 3.48, and 3.48 Å), and using these three reconstructions with the composite map option of REFMAC5^11^, we refined the structure of ID^Ub^ bound to 28 bp of dsDNA at 3.48 Å resolution (Fig. 2a to c; density shown in Extended Data Figs 9c to e, and refinement statistics in Extended Data Table 1). The data sets with 5’ flap DNA, dsDNA and ICL DNA data sets respectively have 98,750, 85,078 and 28,519 particles and their reconstructions extend to 3.8, 3.8 and 4.4 Å (Extended Data Fig. 10a). Within the limits of their resolution, these maps are indistinguishable from that of the nicked DNA (Extended Data Figs 10b to d).

**Figure 2.**
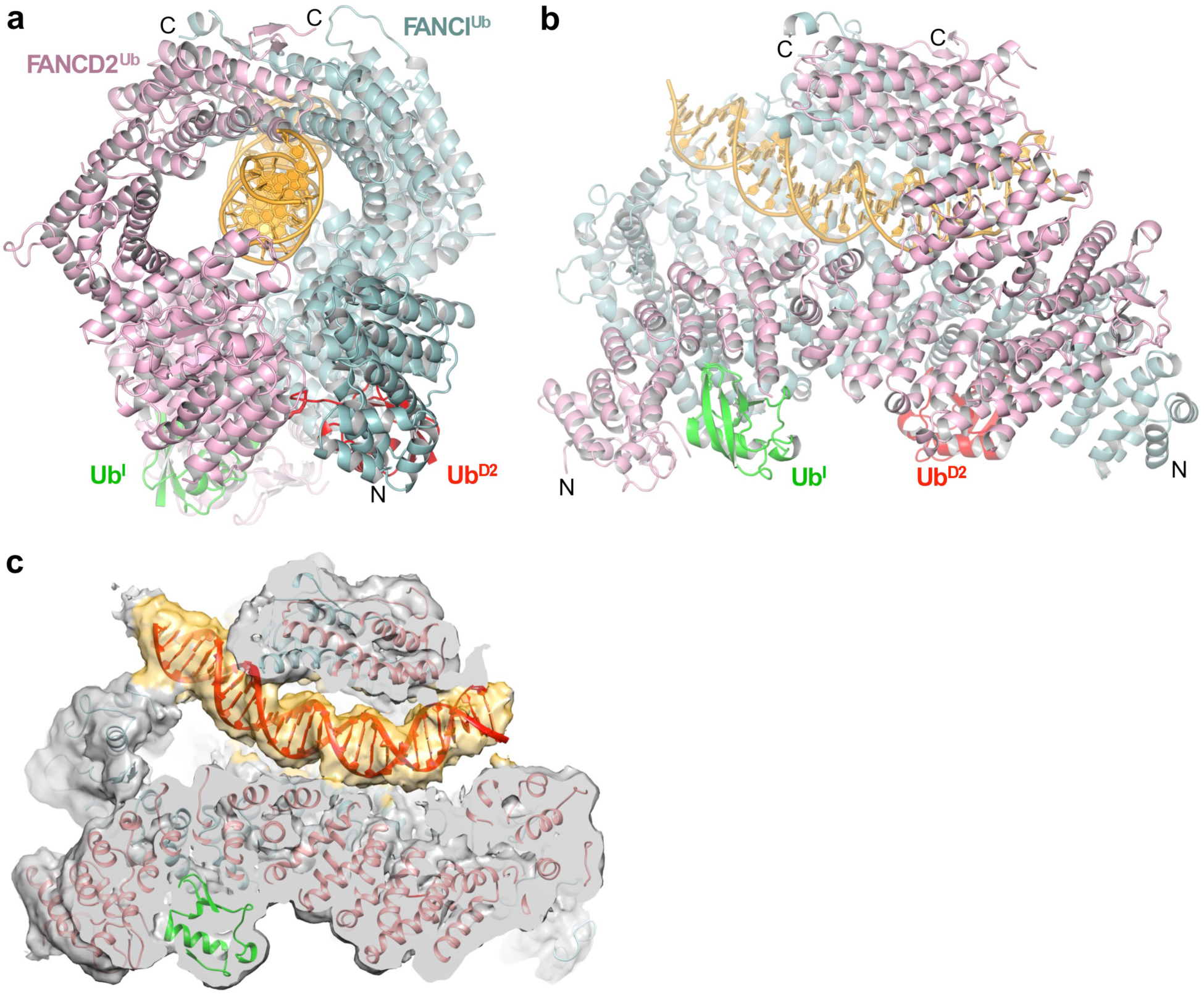
Overall Structure of the mono-ubiquitinated human ID^Ub^ complex bound to nicked dsDNA. **a**, Cartoon representation of the ID^Ub^-DNA complex. FANCD2^Ub^ is pink, FANCI^Ub^ cyan, FANCI ubiquitin (Ub^I^) green, FANCD2 ubiquitin (Ub^D2^) red, and DNA yellow. The N and C termini are labeled, except for the FANCD2 N-terminus that is obscured in this view. FANCI is oriented as in Fig. 1b. **b**, View looking down the left side of the horizontal axis of **a. c**, Cryo-EM density prior to post-processing. Orientation as in **b**. DNA is red, and its density yellow. A portion of FANCD2 above the plane of the figure is cropped to reveal the internal DNA.

Mono-ubiquitination induces a new mode of FANCI-FANCD2 association that results in the conversion of the open-trough structure to a closed ring with the DNA inside (Figs 2a to c). Central to the new mode of heterodimerization are the two ubiquitin molecules, whereby the ubiquitin covalently attached to one paralog binds to the other paralog in a reciprocal fashion.

In the non-ubiquitinated ID complex, the two proteins interact along the length of their NTD solenoids in an antiparallel direction, forming an extended interface that buries a surface area of ∼2500 Å^2^. The interface is continuous except for two narrow openings wherein the ubiquitination sites (Lys523 and Lys561, respectively of FANCI and FANCD2) are embedded (Fig. 1a; Extended Data Figs 2a and b). On ubiquitination, this interface opens up through a relative rotation of FANCI and FANCD2 by 59° and translation by 15 Å about an axis that runs through the NTD-NTD interface (Fig. 3a and b). None of the intermolecular contacts of the non-ubiquitinated interface are retained (Extended Data Fig. 11a). Some of the ID interface residues are repurposed for interactions with the ubiquitin of the other paralog, with the reciprocal interactions contributing ∼1700 Å^2^ of buried surface area (Fig. 3c; Extended Data Fig. 11a). Other residues are repurposed for new intermolecular contacts in a smaller interface between the N-terminal portion of the FANCD2^Ub^ NTD and the C-terminal portion of FANCI^Ub^ CTD (∼1350 Å^2^ buried surface area; Extended Data Fig. 11a).

**Figure 3.**
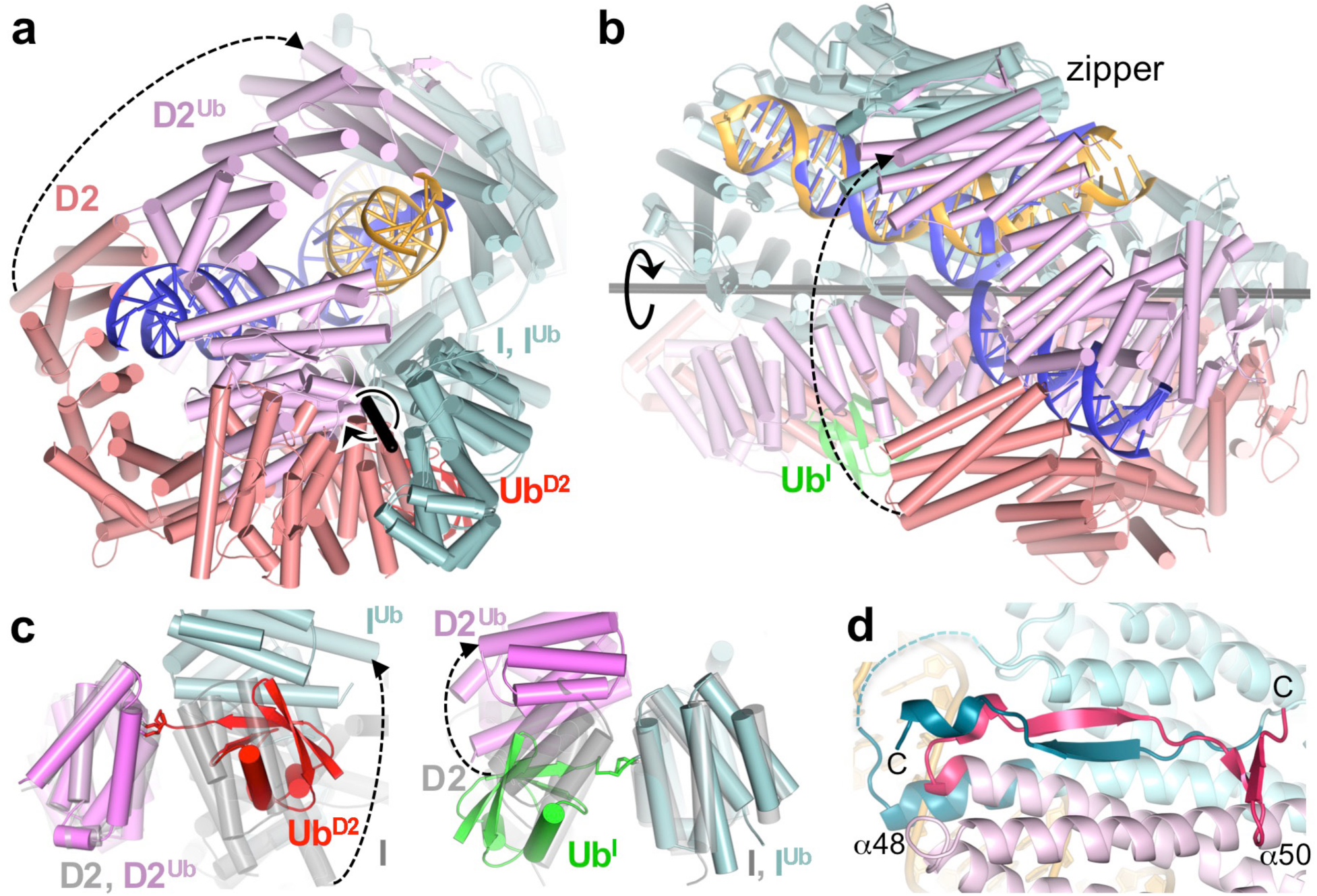
Conformational changes on ubiquitination. **a**, The non-ubiquitinated ID complex (cyan, salmon and blue for FANCI, FANCD2 and DNA, respectively) superimposed on ID^Ub^ (cyan, magenta and yellow for FANCI^Ub^, FANCD2^Ub^, and DNA, respectively) by aligning FANCI. Ub^D2^ is red (Ub^I^ obstructed in this view). The FANCI labels are abbreviated as “I” and those of FANCD2 as “D2”. The rotation axis is shown as a thick black stick going into the plane of the figure with the rotation indicated by a circular arrow. The relative movement of FANCD2 is indicated by a dashed arrow that begins and ends at equivalent structural elements. Orientation similar to Fig. 2a. **b**, View looking down the horizontal axis of **a**. The CTD-CTD interaction region is labeled “zipper”. **c**, Close-up view of the ID and ID^Ub^ complexes superimposed on the FANCD2 ubiquitination site (left) or on the FANCI ubiquitination site (right). The non-ubiquitinated ID complex is transparent and colored gray for all proteins, ID^Ub^ is colored as in **a. d**, Close-up view of the CTD zipper of the ID^Ub^ complex in an orientation similar to **b**. The C-terminal extensions that are unstructured in the non-ubiquitinated complex are shown in darker colors (teal for otherwise cyan FANCI, and hotpink for pink FANCD2). The C-termini of the structures, α48 (FANCI^Ub^) and α50 (FANCD2^Ub^) are labeled. Dashed cyan line is an unstructured loop.

The relative rotation of the two proteins also brings their CTD domains into contact, thereby portions of the previously unstructured C-terminal extensions of FANCI and FANCD2 become structured in a zipper-like interaction (FANCI residues 1233 to 1246 and 1281 to 1297, and FANCD2 residues 1376 to 1400; Figs 3b and b). The zipper involves the FANCI^Ub^ α48 helix forming a coiled coil with α50 of the newly juxtaposed FANCD2^Ub^ CTD, as well as the inter-digitation of the two chains to form an intermolecular β sheet, capped by a short helix in FANCI^Ub^ and a β hairpin in FANCD2^Ub^ (Fig. 3d). The FANCI and FANCD2 re-structured segments end just before their essential EDGE motifs^21^ (residues 1300 to 1303, and 1427-1430, respectively), which remain accessible for possible recruitment interactions with downstream proteins. The zipper buries an additional ∼2200 Å^2^ of surface area, completing the transformation of the previously trough-like structure to a closed ring. The functional importance of ring closing is underscored by the Fanconi Anemia R1285Q mutation^18,22^ in FANCI. Arg1285, which is on the zipper β sheet, forms an intermolecular salt bridge with Glu1365 of FANCD2 in a buried environment, whereas it is unstructured in the non-ubiquitinated ID (Extended Data Fig. 9e).

The conformation of FANCI does not change significantly, and the entire molecule can be superimposed with a 1.6 Å root mean square deviation (r.m.s.d.) in the positions of 1132 C_α_ atoms. By contrast, FANCD2 undergoes two conformational changes important in the remodeling of the complex: the N-terminal part of the NTD rotates by 38° towards FANCI, which is now farther away, to better embrace Ub^I^ and also to form a new interface with FANCI; and the CTD rotates 20° to form the zipper interface with the FANCI CTD (Extended Data Fig. 11b).

FANCI and FANCD2 use similar concave regions of their NTD solenoids (residues 175 to 377 and 174 to 348, respectively) to bind to ubiquitin (Fig. 4a). Both interfaces completely cover the ubiquitin hydrophobic patch consisting of Leu8, Ile44 and Val70 (Figs 4b and c). Ile44 in various combinations with the other two hydrophobic residues is key to the binding of diverse ubiquitin-binding domains such as UBA, UBZ, UIM and CUE^23^, which have structures unrelated to the ubiquitin-binding helical repeats of FANCI and FANCD2 (Extended Data Figs 11c to f). The FANCD2^Ub^-Ub^I^ interactions extend well beyond the hydrophobic patch, owing to the aforementioned conformational change of the FANCD2^Ub^ NTD resulting in a horseshoe-like structure that embraces half the circumference of the ubiquitin, compared to FANCI^Ub^ encircling only a quarter of it (Figs 4a to c; Extended Data Fig. 11b).

**Figure 4.**
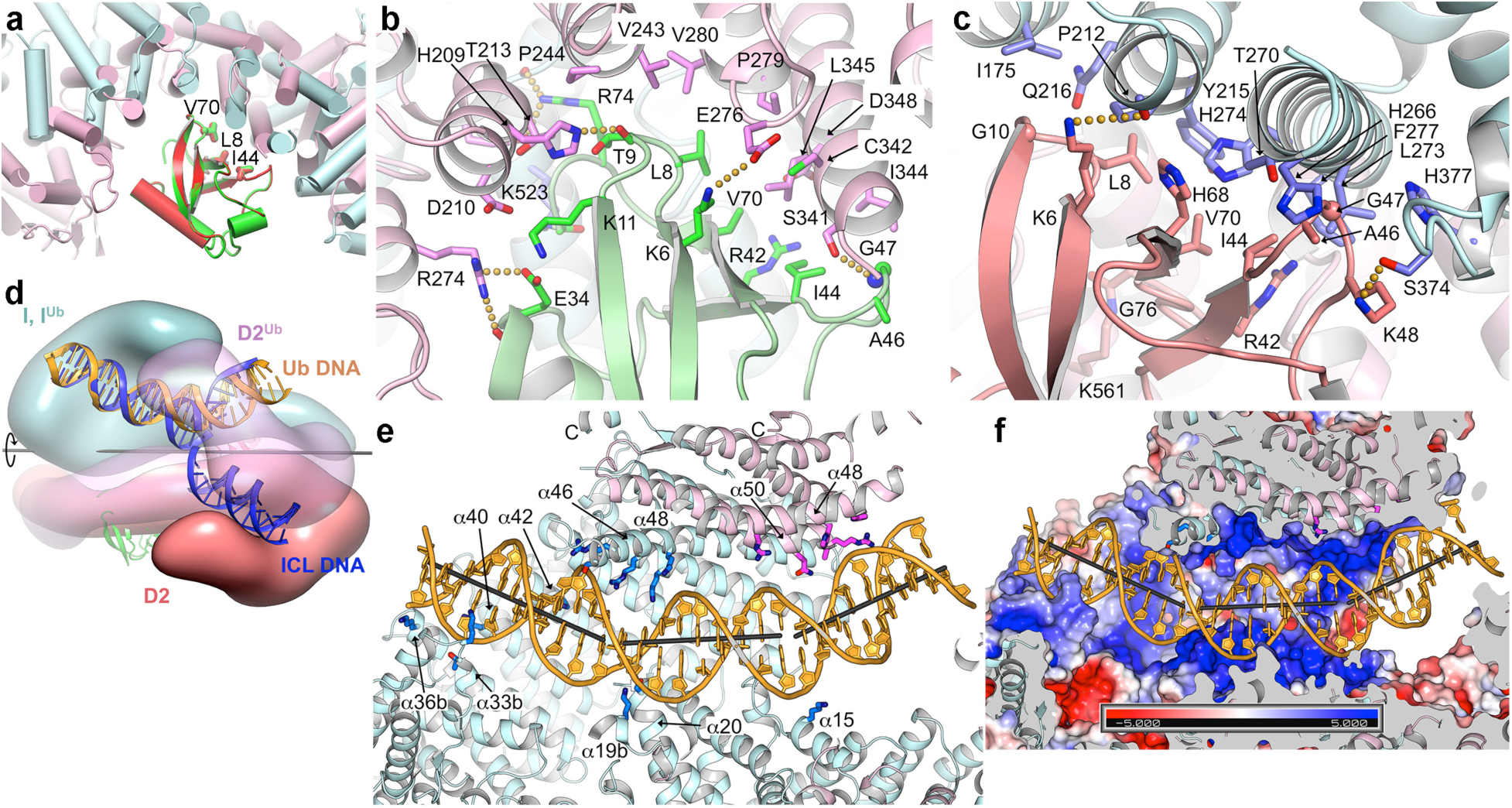
Binding to ubiquitin and to DNA. **a**, Superposition of the non-covalent FANCD2^Ub^-Ub^I^ and FANCI^Ub^-Ub^D2^ interfaces by aligning Ub^D2^ (red) with Ub^I^ (green). The FANC proteins that each ubiquitin is attached to are not shown for clarity (they would be far bellow the plane of the figure). The ubiquitin hydrophobic patch residues (Leu8, Ile44, and Val70) are shown in stick representation for both ubiquitin molecules. Colored as in Fig. 2a. **b**, Close-up view of the non-covalent interface between FANCD2^Ub^ (pink) and Ub^I^ (green) showing the residues within interaction distance as sticks (darker colors). Yellow dotted lines indicate potential hydrogen bonds. **c**, Close-up view of the reciprocal non-covalent interface between FANCI^Ub^ (cyan) and Ub^D2^ (green) as in **b. d**, Rendering of ID^Ub^-DNA and ID-ICL DNA superimposed on their respective FANCI proteins. Colored as in Fig. 2a. FANCD2^Ub^ is semi-transparent to reveal the DNA inside. **e**, Cartoon representation showing ID^Ub^ side chains within contact distance of the DNA as sticks. The structural elements involved and the C-termini are labeled. Colored as in Fig. 2a except for darker side chains. Gray sticks show the helical axes for the three relatively straight segments of the DNA. The majority of FANCD2^Ub^ and parts of FANCI^Ub^ are clipped above the plane of the figure to make the DNA visible. Shown are FANCI^Ub^ side chains for Lys291 (α15), Lys397 (α19b), Ser411 (α20), Lys793, Thr794 on α33b, Lys898 (α36b), Lys980 (α40), Lys1026 (α42), Lys1164 (α46), and Thr1238, Arg1242, Arg1245 and Lys1248 on the extended α48; FANCD2^Ub^ side chains for His1288, His1292, and Arg1299 on α48, and Thr1351, Arg1352 and Gln1355 on α50. **f**, Same view as in **e** showing the molecular surface colored according to electrostatic potential calculated with PYMOL in the absence of DNA (colored −5 to +5 kT blue to red). The molecular surface interior is uniformly gray.

It has been suggested that one function of ID mono-ubiquitination may be the recruitment of downstream effectors that contain ubiquitin-binding domains^1,24^. However, the sequestration of the entire ubiquitin hydrophobic patch on both Ub^I^ and Ub^D2^ indicates they are unlikely to play this role, at least with cofactors containing the aforementioned ubiquitin-binding domains. This is consistent with a recent report that the UBZ-containing FANCP nuclease scaffold does not require FANCD2 ubiquitination for recruitment to ICL sites^25^.

The mono-ubiquitination induced conformational change results in the remodeling of the bound DNA, converting the non-collinear arrangement of the FANCI- and FANCD2-associated DNA duplexes to a continuous, though bent duplex (Figs 3b and 4d). Two bends, of 26° and 31°, are centered on the 11^th^ and 21^st^ base pairs from the FANCI end, respectively, with large roll values over three base-pair steps.

DNA binding by the ID^Ub^ complex involves structural elements that mostly overlap with those of the ID complex, but there are differences associated with the new ID and DNA conformations. As with the ID-ICL DNA complex, the FANCI^Ub^ groove partially encircles the DNA through basic and polar residues from α33b and α36b on one side of the groove and α40, α42 on the other (Figs 4e and f). Near the middle of the DNA, however, the extension of α48 that results from coiling with FANCD2^Ub^ α50 (Fig. 3d) gives rise to a new semicircular basic groove, between α46 and α48 on one side and the NTD α19b and α20 on the other, into which dsDNA binds (Figs 4e and f). This is associated with one of the two DNA bends, which redirects the duplex away from clashing with the new position of FANCD2 (Figs 4e and f). Thereafter, the second bend occurs as the dsDNA is redirected by the FANCD2 CTD, which uses the localized patch of α48 and α50 to bind to the duplex analogously to the ID-ICL DNA complex. However, because of the new position of the FANCD2 CTD, the FANCD2^Ub^-associated dsDNA coincides with the location of ssDNA in the ID-ICL DNA complex, and displaces it (Fig. 4d).

We have not been able to locate the nick of the nicked DNA in the map, which looks indistinguishable from the 3.8 Å map of ID^Ub^ bound to canonical dsDNA (Extended Data Fig. 10c). While this could be due to the inadequate resolution of the nicked DNA density, the maps of ID^Ub^ bound to 5’ flap DNA or ICL have no trace of their ssDNA branches neither (Extended Data Figs 10b and d). This suggests either that ID^Ub^ binds to these substrates in multiple registers without a preferred location for the nick or flap, or it is just binding to the canonical dsDNA arms, which are 29 bp in the nicked and 5’ flap substrates and 21 bp in ICL DNA. Either case implies that the ID^Ub^ complex has lost the specificity the non-ubiquitinated ID complex has for branched DNA structures. Accordingly, the dissociation constants of ID^Ub^ for dsDNA, nicked, 5’ flap, ICL and fork DNA vary by less that a factor of 2, with dsDNA and nicked DNA exhibiting slightly tighter *K*_d_ values the rest (27, 20, 32, 33, and 30 nM, respectively, Extended Data Fig. 10e).

Our data reveal the function of ID mono-ubiquitination is to completely re-model FANCI-FANCD2 association, expanding our understanding of the roles of mono-ubiquitination. For the ID complex, the functional significance of this re-modeling is to convert it to a clamp that could slide away from its initial location at the ICL DNA. In principle, this would allow downstream nucleases and other factors to act on the ICL, with the ID^Ub^ clamp coordinating the repair reactions, serving as a processivity factor, or protecting the dsDNA.

## Acknowledgments

We thank the staff of the MSKCC Cryo-EM facility, the NYSBC Simons Electron Microscopy Center, and the HHMI Cryo-EM facility for help with data collection. Supported by HHMI and National Institutes of Health grant CA008748.

## Author Contributions

R.W. carried out the biochemical experiments and collected and analyzed the cryo-EM data, S.W. prepared the FA Core complex, R.W. and S.W. carried out the ubiquitination reactions, A.D. collected cryo-EM data and carried out biochemical experiments, C.P. carried out biochemical experiments, and N.P.P. analyzed the data and wrote the manuscript.

## Author Information

The authors declare no competing financial interests. Correspondence and requests for materials should be addressed to N.P.P.

## METHODS

### Protein Expression and purification

Full length human FANCI with a non-cleavable C-terminal His_6_ tag and full length human FANCD2 with an N-terminal His_6_ tag followed by a TEV protease cleavage site were co-expressed in Hi5 insect cells using baculovirus. Cells were lysed in 50 mM Tris-HCl, 200 mM NaCl, 5 % (v/v) glycerol, 0.5 mM TCEP, pH 8.0, and protease inhibitors. After Ni^2+^ affinity chromatography and overnight cleavage of the His_6_ tag by TEV protease, the FANCI and FANCD2 proteins were purified by ion exchange (MonoQ) chromatography, which dissociated the two proteins. The FANCI and FANCD2 proteins were then combined at a 1:1 molar ratio, concentrated by ultrafiltration to ∼20 mg/ml, and the ID complex was purified by gel filtration chromatography in 20 mM Tris-HCl, 150 mM NaCl, 0.5 mM TCEP, pH 8.0. For the FA Core complex, Flag-tagged FANC A, B, C, E, F, G, L, and FAAP100 were cloned into three modified pCDNA3.1 plasmids with different drug resistance genes, and were used to transfect HEK 293F cells. A stably-transfected cell line with the highest expression of all eight subunits was then adapted for growth in suspension. The FA Core complex was purified using anti-Flag M2 agarose beads (Sigma), ion-exchange (MonoQ) and gel-filtration chromatography (Superose 6) in 20 mM Bicine-HCl, 150 mM NaCl, 0.1 mM TCEP, pH 8.0.

### In vitro ubiquitination

Ubiquitination reactions, carried out in 20 mM Tris-HCl, 150 mM NaCl, 1 mM DTT, pH 8.0, contained 40 µM human Flag-tagged ubiquitin (Fisher Scientific U-120), 0.5 µM human His_6_-tagged Ubiquitin E1 enzyme (Fisher Scientific, E304050), 2.4 µM human His_6_-UBE2T E2 enzyme (Fisher Scientific, E2-695), 5 mM adenosine triphosphate, 3 µM core complex, 10 µM ID complex and 30 µM DNA. 60 µL reactions were set up on ice and incubated at 28°C for 1-2 hours. Samples were separated by SDS-PAGE with NuPAGE 3%–8% Tris-Acetate gels (Invitrogen) and the mono-ubiquitinated FANCI and FANCD2 were verified using mass spectrometry.

### Cryo-EM sample preparation and data collection

For the non-ubiquitinated ID complex bound to DNA, the ID peak from the gel filtration chromatography was combined with a 3-fold molar excess of DNA, and the mixture was diluted to 2 mg/ml for the ICL DNA sample and to 3 mg/ml for the other DNA substrates, in 20 mM Tris-HCl, 150 mM NaCl, 0.5 mM TCEP, pH 8.0. The sample (3 µl) was applied to glow discharged UltrAuFoil 300 mesh R1.2/1.3 grids (Quantifoil). Grids were blotted for 1.5 s at 16° C or 22° C and ∼100 % humidity and plunge-frozen in liquid ethane using a FEI Vitrobot Mark IV. For the mono-ubiquitinated ID^Ub^ complex, the in vitro ubiquitination reaction described above was fractionated by gel-filtration chromatography (Superose 6) in 20 mM Tris-HCl, 150 mM NaCl, 0.2 mM TCEP, pH 8.0, and the peak corresponding to the ID^Ub^-DNA complex was concentrated by ultrafiltration to ∼1.5 mg/ml. Grids were prepared as with the non-ubiquitinated complex. All data were collected with a Titan Krios microscope operated at 300 kV and Gatan K2 Summit camera. Most data were collected with a 1.089 Å pixel size and 10.0 electrons per pixel per second at the MSKCC Cryo-EM facility, with additional data collected at the NYSBC Simons Electron Microscopy Center (1.09 Å pixel size), and at the HHMI Cryo-EM facility (1.04 Å pixel size and 8.0 electrons per pixel per second).

### Cryo-EM image processing

The super-resolution movies were initially aligned with MOTIONCOR2^26^, and the contrast transfer function (CTF) parameters were estimated with CTFFIND4^27^. All 2D/3D classifications, 3D refinements and other image processing were carried out with RELION-3^9^. For the data from the ID-ICL DNA complex and all the ID^Ub^-DNA complexes, Bayesian beam induced motion correction, scale and B-factors for radiation-damage weighting, and per particle refinement of CTF parameters were also applied^28^. All reported map resolutions are from gold-standard refinement procedures with the FSC=0.143 criterion after post-processing by applying a soft mask. To account for the large conformational changes in the FANCD2 CTD of the ID-ICL DNA complex, we performed multi-body refinement with two soft masks. The larger-body (body1 plot in Extended Data Fig. 1a) mask covered FANCI, the FANCI-bound dsDNA and ssDNA, and residues 45 to 623 of FANCD2, and the smaller-body (body2 in Extended Data Fig. 1a) mask covered FANCD2 residues 624 to 1376 and the FANCD2-associated dsDNA. Multi-body refinement was carried out with signal subtraction outside the masks. The ID^Ub^ complex does not exhibit conformational flexibility in the FANCD2 CTD. The three focused 3D refinements of the ID^Ub^-nicked DNA data improved the resolution only marginally, but there was noticeable improvement in side chain density and continuity. Of the three partially overlapping masks, one mask (focus1 plot in Extended Data Fig. 9a) covered the C-terminal portions of FANCI (residues 598-1298) and FANCD2 (residues 625-1400) and the entire DNA, corresponding roughly to the top half of Fig. 2a. A second mask (focus2 plot in Extended Data Fig. 9a) covered FANCI residues 1-376, FANCD2 residues 447-1400, Ub^D2^, and the FANCD2-proximal half of the DNA, and corresponds roughly to the right half of Fig. 2b. And a third mask (focus3 plot in Extended Data Fig. 9a) covered FANCI residues 288-1297, FANCD2 residues 45-466, Ub^I^, and the FANCI-proximal half of the DNA, roughly the left half of Fig. 2b.

### Cryo-EM structure refinement

Model refinement was done with REFMAC5 modified for cryo-EM^11^. For the ID-ICL DNA complex, the two multi-body refinement maps were combined with the composite sfcalc option of REFMAC5 to construct a single set of structure factors to 3.3 Å, the resolution of the larger body1, and refinement was carried out at this resolution. As the resolution of the smaller body2 map is lower (3.8 Å), this results in higher temperature factors for the portion of the model within this map. The structure factors for the ID^Ub^-nicked DNA complex were calculated by combining the three focused 3D reconstructions similarly, except refinement was carried out at 3.48 Å, the highest resolution common to all 3 maps. The coordinates were assigned to the three focused maps as follows: FANCI residues 960-1297, FANCD2 residues 1150-1400, and the DNA to focus1; FANCI residues 1-326, FANCD2 residues 455-1145, and Ub^D2^ to focus2; FANCI residues 960-1297, FANCD2 residues 1150-1400, and the DNA to focus1; FANCI residues 327-959, FANCD2 residues 45-454, and Ub^I^ to focus3. Both refinements included secondary structure restraints (SSR) generated by ProSMART^11^, and TLS refinement.

The refined ID-ICL DNA model lacks the following unstructured regions: 250-259, 401-408, 551-574, 685-695, 935-948, 1111-1125, 1222-1246, 1281-1328 (C-terminus) of FANCI, and 1-44, 122-129, 313-337, 589-604, 708-725, 852-915, 947-959, 982-1000, 1043-1050, 1146-1149, 1216-1219, 1377-1451 of FANCD2. As discussed in the main text (Fig. 3d), in the refined ID^Ub^-nicked DNA model FANCI residues 1233-1246 (α48 extension) and 1281-1297 (C-terminal residues), and FANCD2 residues 1377-1400 become ordered as part of the zipper interface. The unstructured regions of the non-ubiquitinated ID-ICL DNA complex generally correspond well with the unstructured regions in the mouse ID crystal structure, or the regions deleted from the mouse FANCI and FANCD2 constructs used in those crystallization experiments based on susceptibility to limited proteolysis^7^. The one exception is FANCI residues 551-574, where the corresponding region of mouse FANCI is well ordered within the FANCI-FANCD2 interface. The FANCD2 region with which this segment packs in the mouse ID complex is shifted by ∼5 Å away from FANCI in the human ID complex. This segment also contains phosphorylation sites for ATR kinase^7,29^.

### DNA-binding assays

Electrophoretic mobility shift assays (EMSA) ssays were performed using the 40 to 42 bp (or equivalent) DNA substrates shown in the table below. For the non-ubiquitinated ID complex, the DNA-binding reactions contained unlabeled 20 bp dsDNA as nonspecific competitor. Reactions (15 μl) were assembled by mixing the indicated ^32^P-labeled DNA substrates (0.5 nM) with the unlabeled dsDNA (1.4 μM) and adding the ID complex in 20 mM Tris-HCl, 150 mM NaCl, 5 % Glycerol, 0.1 mg/ml BSA, 0.5 mM TCEP, pH 8.0. They were incubated on ice for 30 min, followed by electrophoresis at 4 °C on 4 % (w/v) polyacrymide gels in 0.5x TBE buffer. The dried gels were quantitated using a phosphorimager, and the data were fit to a one-site cooperative binding model by minimizing the sum of square of the differences. For the mono-ubiquitinated ID^Ub^ complex, the DNA that remained bound to the ID^Ub^ complex during gel-filtration was removed by anion-exchange chromatography. This also dissociated the FANCI^Ub^ and FANCD2^Ub^ proteins, which were then concentrated separately by ultrafiltration. Binding reactions were assembled by first mixing FANCI^Ub^ with the indicated ^32^P-labeled DNA substrate (0.5 nM) and then adding FANCD2^Ub^ at a one molar ratio to FANCI^Ub^. The subsequent steps were performed as with the non-ubiquitinated ID complex. The reactions did not contain the unlabeled dsDNA competitor used with the non-ubiquitinated ID complex.

### DNA substrates

Substrates for ID and ID^Ub^ DNA complexes were prepared by annealing the oligonucleotides listed below. The ICL DNA was prepared as described^8^. Briefly, we used the Cu(I)-catalyzed azide-alkyne cycloaddition between an N4-(3-azidopropyl) modified cytosine on a dCdG step of one DNA strand, and an N4-propargyl modified cytosine on the complementary strand, crosslinking the N4 positions of the two cytosines with a triazole moiety. The A3 and A4 oligonucleotides were synthesized by Sigma incorporating N4-(3-azidopropyl) deoxycytidine and N4-propargyl deoxycytidine, respectively, at the positions indicated by the bold “**C**”. For the A3 oligonucleotide, the synthesis involved first adding an N4-chloropropyl deoxycytidine phosphoramidite to avoid an azide-phosphoramidite side reaction, with the chloropropyl group subsequently converted to azidopropyl on the beads. The oligonucleotides were HPLC purified by Sigma. For the crosslinking, the two oligonucleotides, each at 0.2 mM concentration, were incubated with 2 mM CuSO_4_, 10 mM Sodium Ascorbate and 1 mM Tris(3-hydroxypropyltriazolylmethyl)amine in 10 mM HEPES-Na, 50 mM NaCl, pH 7.5, for 30 minutes at room temperature. The reaction products were separated by denaturing PAGE (12% polyacrylamide, 8 M urea), and the crosslinked product was isolated from the gel slice by electroelution (Whatman Elutrap). The ICL was confirmed by liquid chromatography coupled electrospray ionization mass spectroscopy, performed by Novatia (Newtown, PA).

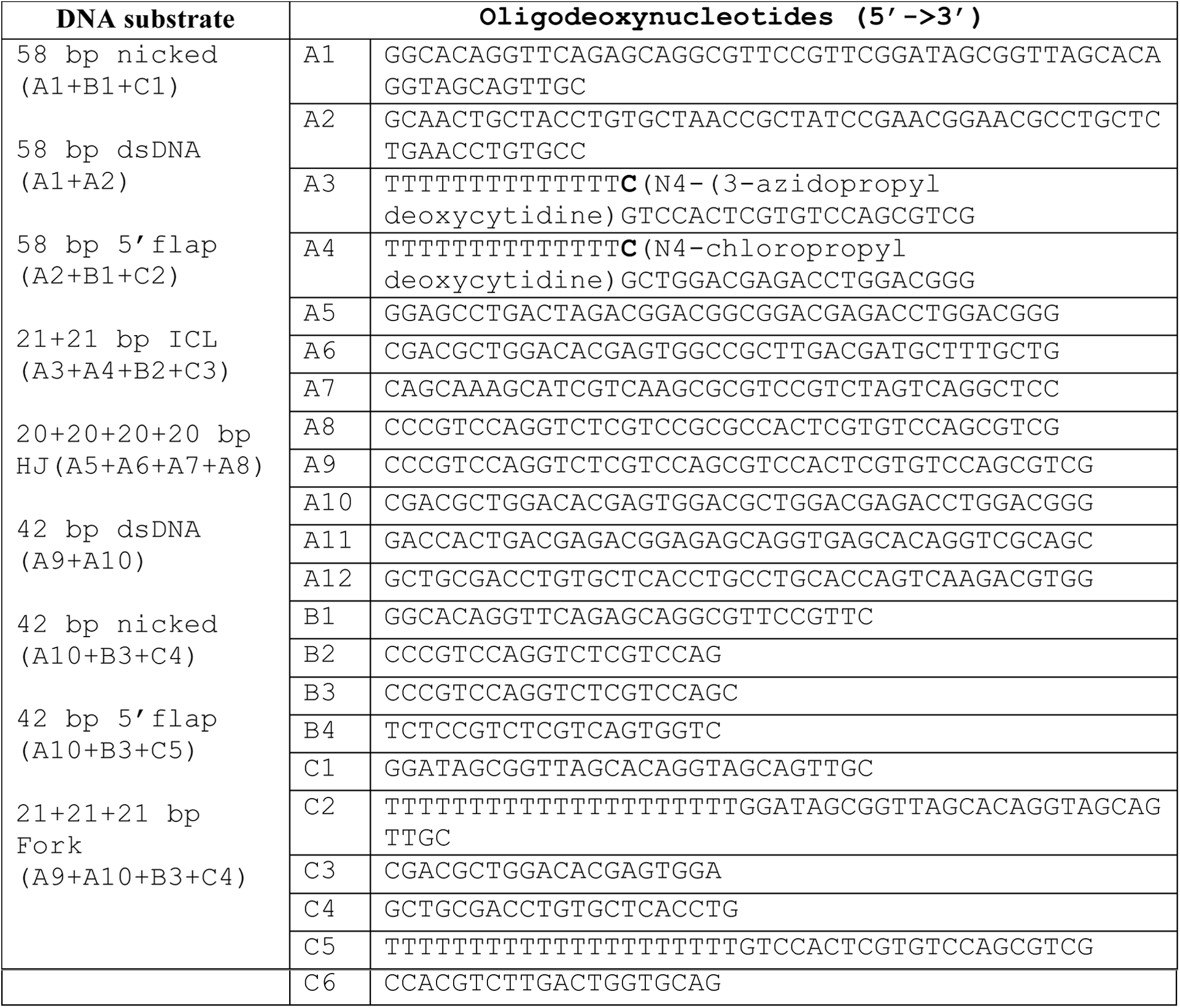

### Data availability

The ID-ICL DNA and ID^Ub^-DNA coordinates and the corresponding cryo-EM maps, including the focused reconstructions and the composite map used in refinement, are being deposited with the Protein Data Bank and the Electron Microscopy Data Bank, respectively.

