## Supplementary material for "DNA clamp function of the mono-ubiquitinated Fanconi Anemia FANCI-FANCD2 complex": SupplemntaryMaterials

**EXTENDED DATA**

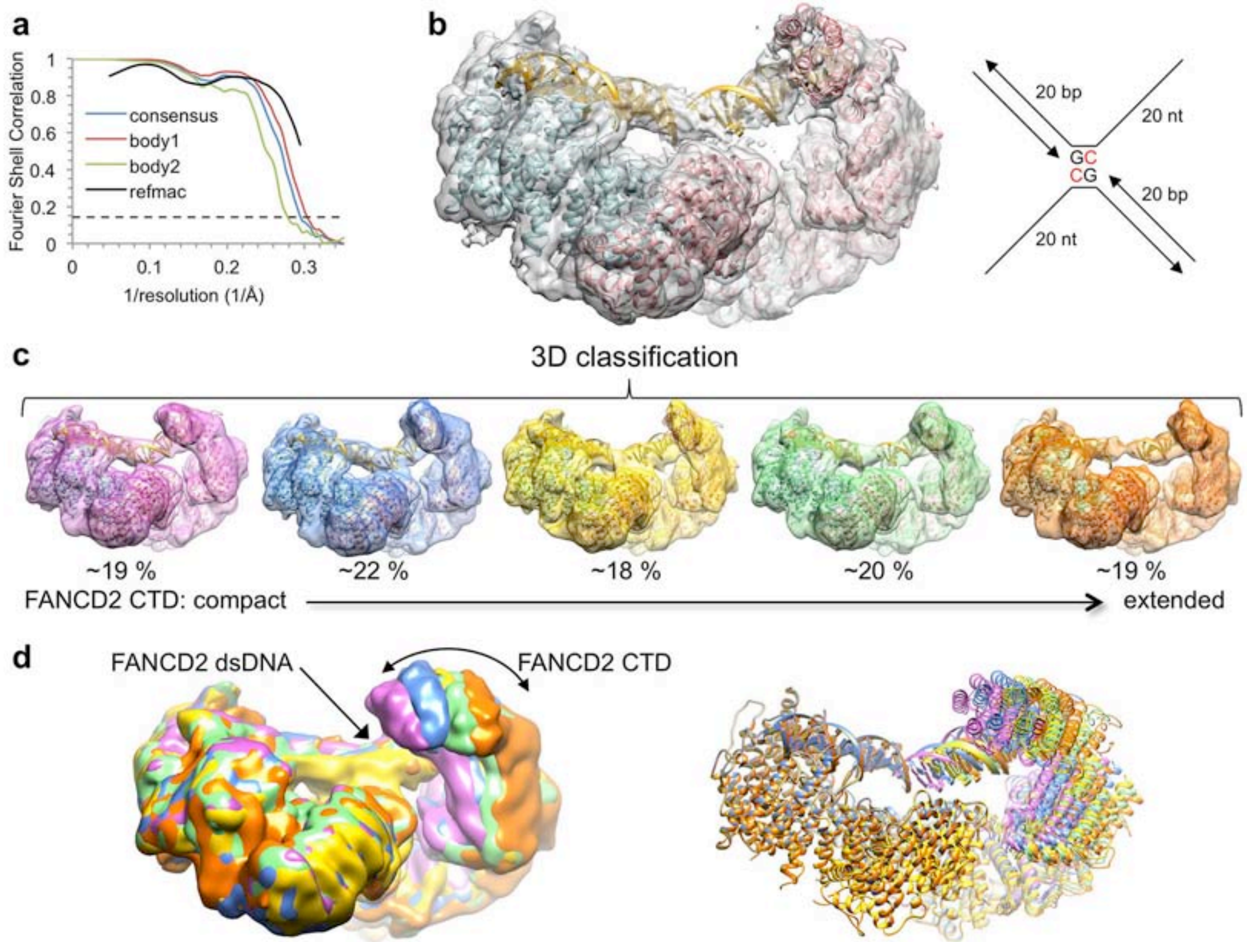


**Extended Data Figure 1 | Cryo-EM reconstruction of the non-ubiquitinated ID complex bound to ICL DNA.**

**a,** Graph shows gold-standard FSC plots between two independently refined half-maps for the consensus reconstruction (blue curve), the RELION3 multi-body refinement of the larger body1 consisting of FANCI, FANCI-bound dsDNA and ssDNA, and FANCD2 residues 43-623 (red curve), and for the smaller body2 consisting of FANCD2 residues 624-1376 and the FANCD2-associated dsDNA (green curve). The FSC curve for the final model versus the composite map combining the cryo-EM maps of the two bodies from REFMAC5 is shown in black. Dashed line marks the FSC cutoff of 0.143.

**b,** Cryo-EM map from the consensus reconstruction prior to post processing by RELION3 (before B-factor sharpening) with the cartoon representation of the refined model superimposed (FANCI in cyan, FANCD2 in pink, and DNA in gold). Schematic of the ICL DNA is shown to the right of the map, with the deoxycytidine bases that are crosslinked by a triazole colored red. The 20 nt ssDNA arms consist of $(dT)_{20}$ to minimize secondary structure.

**c,** 3D classification of the particles showing the conformational flexibility of FANCD2. The 3D classes are arranged starting with the most compact conformation where the FANCD2 CTD is closer to its NTD. Also shown is the refined consensus model rigid-body fitted into each class and colored as in **a**.

**d,** The five 3D classes are superimposed by aligning the FANCI-portion of each map (left), or of each pdb (right) colored as in **c**.

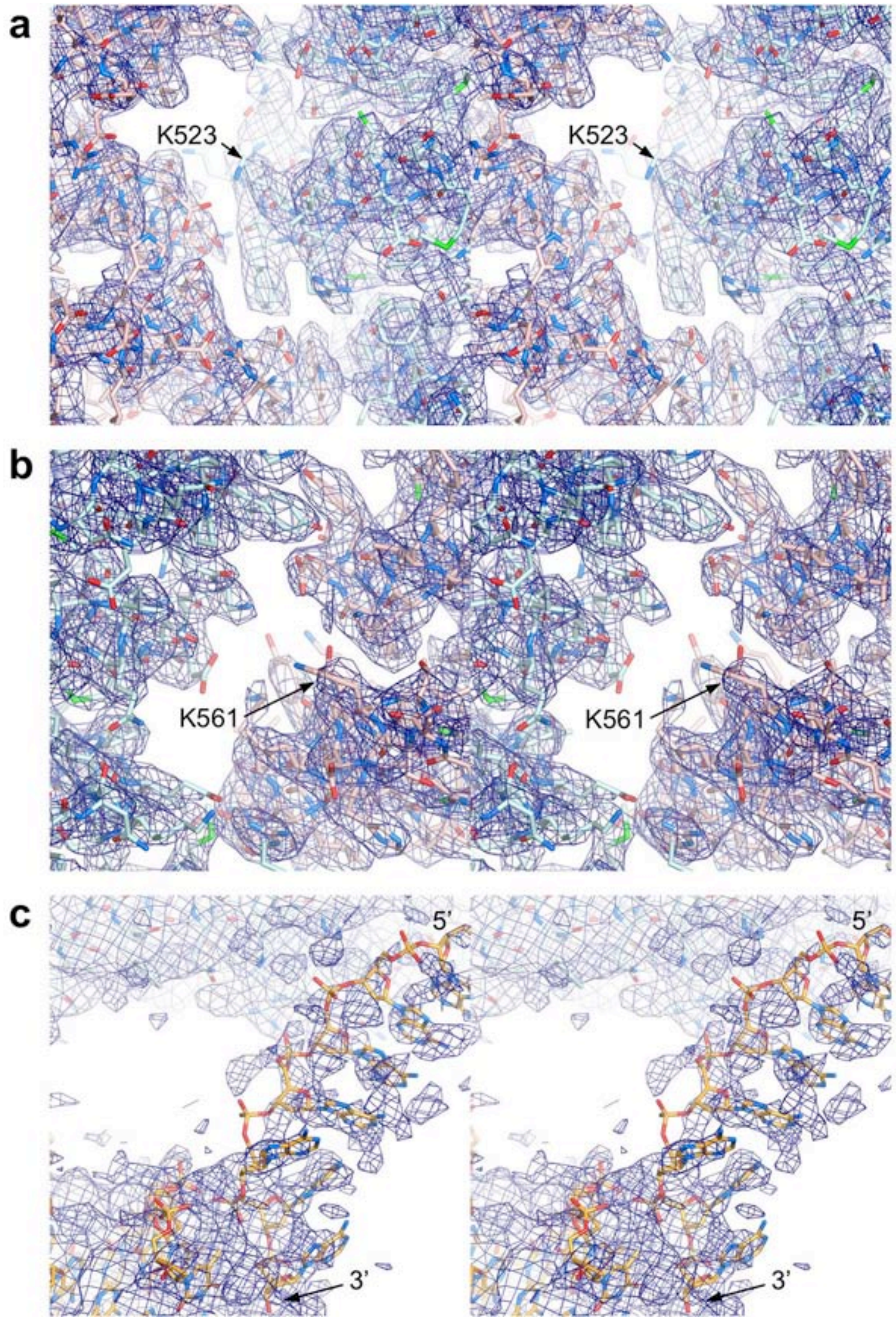


**Extended Data Figure 2 | Cryo-EM density from post-processed maps of non-ubiquitinated ID complex bound to ICL DNA.**

**a,** Stereo view of the 3.3 Å cryo-EM density of the post-processed reconstruction using multi-body refinement. Map shows the vicinity of the FANCI ubiquitination site (Lys523 marked) with portions of FANCI residues 475-593 (cyan) and FANCD2 residues 174-287 (pink) shown in stick representation. O and N atoms are colored half-bonded red and blue, respectively, for both proteins.

**b,** Stereo view of the map from **a** showing the vicinity of the FANCD2 ubiquitination site (Lys561 marked), with portions of FANCD2 residues 482-578 and of FANCI residues 123-223 shown as in **a**.

**c,** Stereo view of the map from **a** focusing on the ssDNA (5' and 3' ends marked) at the FANCI CTD, as well as a portion of the FANCI-bound dsDNA. The DNA is in stick representation colored half-bonded yellow, red and blue for C, O, N atoms, respectively. The map is shown at a low contour level because the ssDNA has high temperature factors, and its density is broken up due to the B-factor applied in post-processing being calculated from the entire map. ssDNA density before post-processing can be seen in the panel of maps in Extended Data Fig. 3.

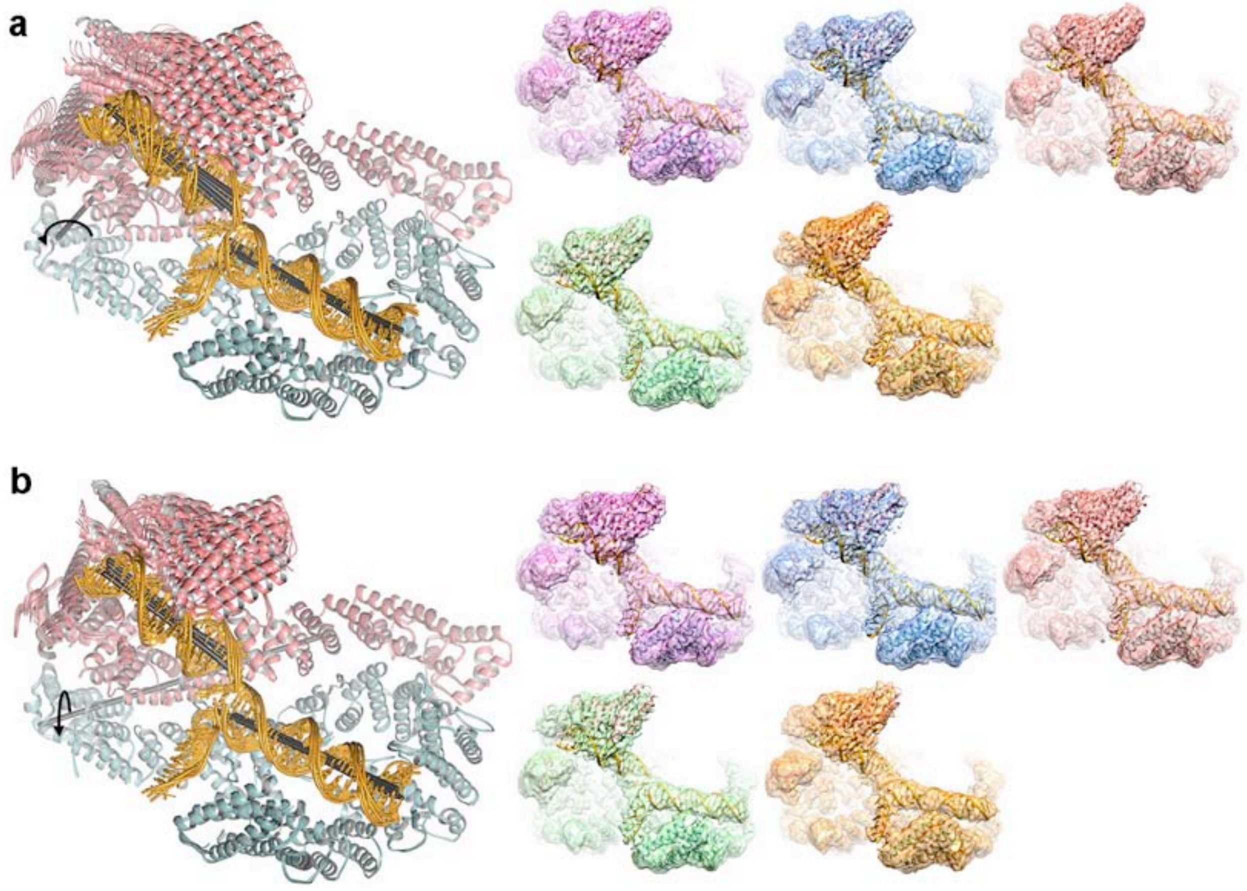


**Extended Data Figure 3 | FANCD2 CTD conformations from the principle component analysis of multi-body angles.**

**a,** Cryo-EM reconstructions using particles from the top component of the principle component analysis (PCA) of the multi-body refinement angles. This component accounts for 21.5 % of the variance in the relative orientation of the FANCD2 CTD (Supplementary Video 1). Left, five ID-ICL DNA models refined in real-space with PHENIX (overall solvent-corrected resolution ranging from 3.67 to 3.87 Å) against maps reconstructed with particles derived from five bins of eigenvalues for the top eigenvector. This PCA component corresponds to a rotation of up to 16° (curved arrow) about an axis running through the HD domain roughly perpendicular to the plane of the figure (gray stick). The helical axes of the individual duplexes are shown as black sticks. Right, the corresponding maps without post-processing, colored sequentially as in Extended Data

Fig. 1a starting with the conformation (pink map) where the FANCD2 CTD is closest to its NTD.

**b,** The second component from the PCA analysis accounts for 17% of the variance in the relative orientation of the FANCD2 CTD (Supplementary Video 2). It corresponds to an up to ~10° rotation (curved arrow) about an axis (gray stick) roughly parallel to the plane of the figure. Left are the refined models, and right the maps as in **a**.

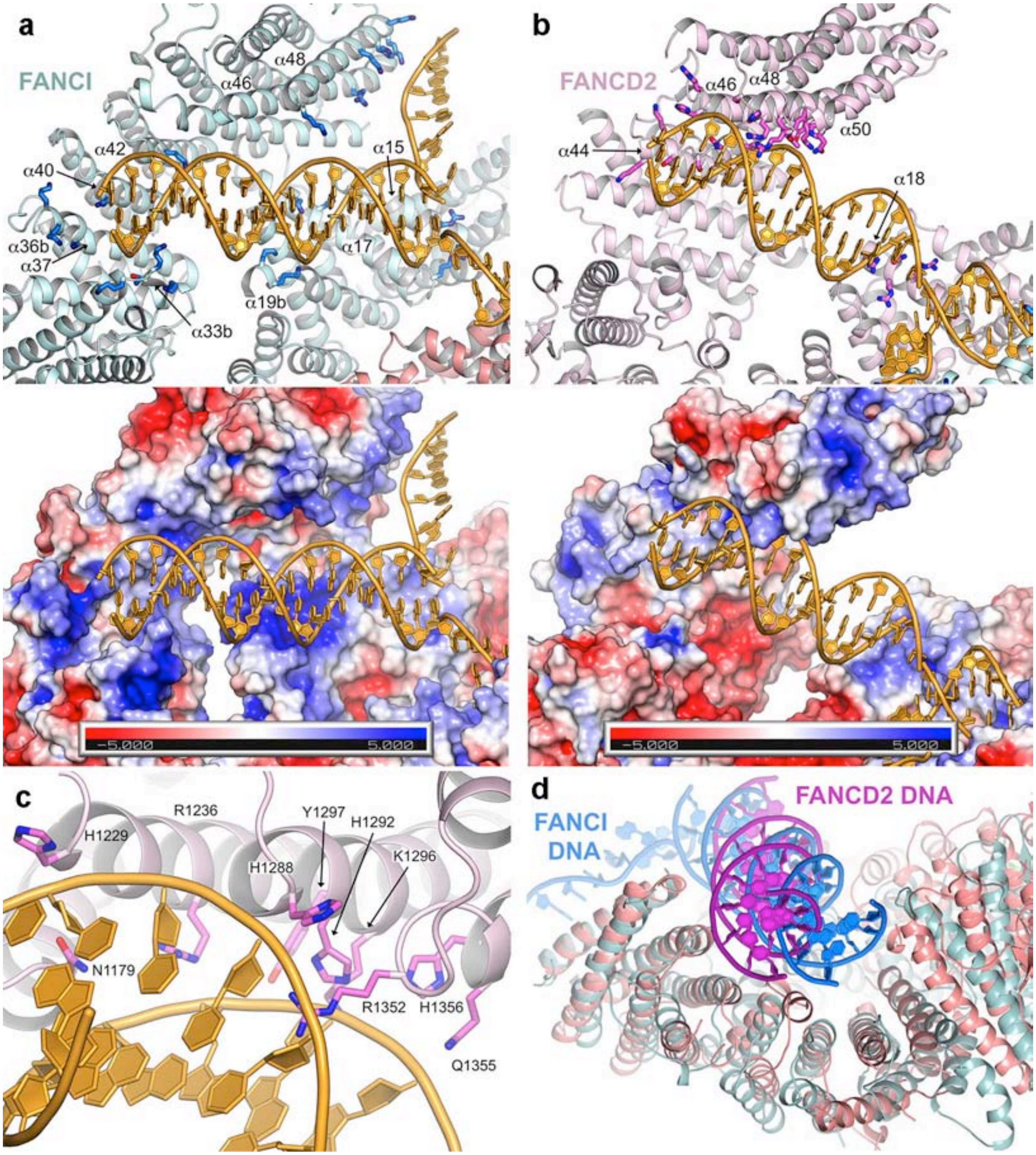


**Extended Data Figure 4 | DNA binding by the non-ubiquitinated ID complex.**

**a,** DNA binds to an extended basic surface of FANCI. Cartoon representation showing FANCI side chains within contact distance of the DNA (top), and molecular surface colored according to the electrostatic potential calculated with PYMOL (bottom, colored -5 to +5 kT blue to red). The end of the dsDNA binds to a semi-circular groove consisting of helices $\alpha$33b, $\alpha$36b, $\alpha$37, $\alpha$40 and $\alpha$42 (secondary structure elements numbered as in the mouse ID structure[30], with insertions denoted by letters after the helix number). This is analogous to the 7.8 Å crystallographic map of mouse FANCI bound to Y DNA, with the N-termini of helices and inter-helix loops providing multiple basic residues. The ICL-proximal portion of the duplex, which is absent from the shorter DNA used in the mouse FANCI crystals, is positioned against basic residues emanating from helices $\alpha$15, $\alpha$17 and $\alpha$19b. The ssDNA rests against the sides of the $\alpha$48 and $\alpha$49 helices. The overall DNA density is of lower resolution than the surrounding protein, and in the refined model the DNA has high temperature factors suggesting it is significantly more mobile than the surrounding protein elements. Figure shows side chains for Arg287 on $\alpha$15, Arg321, Lys336 and Lys339 on $\alpha$17, Lys396 and Lys397 on $\alpha$19b, Lys791, Lys793, Thr794 and Lys795 on $\alpha$33b, Lys897, Lys898 and Lys902 on $\alpha$36b, Lys980 on $\alpha$40, Lys1026 on $\alpha$42, Arg1178 on $\alpha$46, and Lys1262, His1266, Lys1269 and Lys1270 on $\alpha$48.

**b,** FANCD2-DNA contacts are localized to the last four helical repeats of the CTD and a small patch of basic residues on the NTD. Top figure shows the residues within contact distance of the DNA. The CTD residues involve the N-terminal portions of the inner helices of the helical repeats: Lys1172, Lys1174, Ser1175, Ser1178, Asn1179 and His1183 on $\alpha$44, Arg1128, His1229 and Arg1236 on $\alpha$46, Ser1287, His1288, His1292, Lys1296 and Tyr1297 on $\alpha$48, and Thr1351, Arg1352, Gln1355 and His1356 on $\alpha$50. NTD residues on $\alpha$50 are Arg401, Arg404, Asn405 and Arg408. Bottom figure shows the corresponding molecular surface colored according to the electrostatic potential calculated with PYMOL (bottom, colored -5 to +5 kT blue to red). Note the absence of a basic patch at the HD-portion of the semi-circular groove (lower left portion of figure) compared to that of FANCI in **a**.

**c,** Close-up view of the FANCD2 Arg1352 side chain inserting into the minor groove of the DNA, and the residues that contact the flanking phosphodiester backbone.

**d,** Superposition of the DNA-binding region of FANCD2 (pink) CTD on the corresponding region of the FANCI paralog (cyan) showing the different orientations of the FANCI dsDNA (blue) and FANCD2 dsDNA (magenta) in the semi-circular grooves of the respective proteins. Residues 905-1269 of FANCD2 were aligned on residues 1058-1376 of FANCI with a 2.2 Å r.m.s.d. in the positions of 185 $C_\alpha$ atoms.

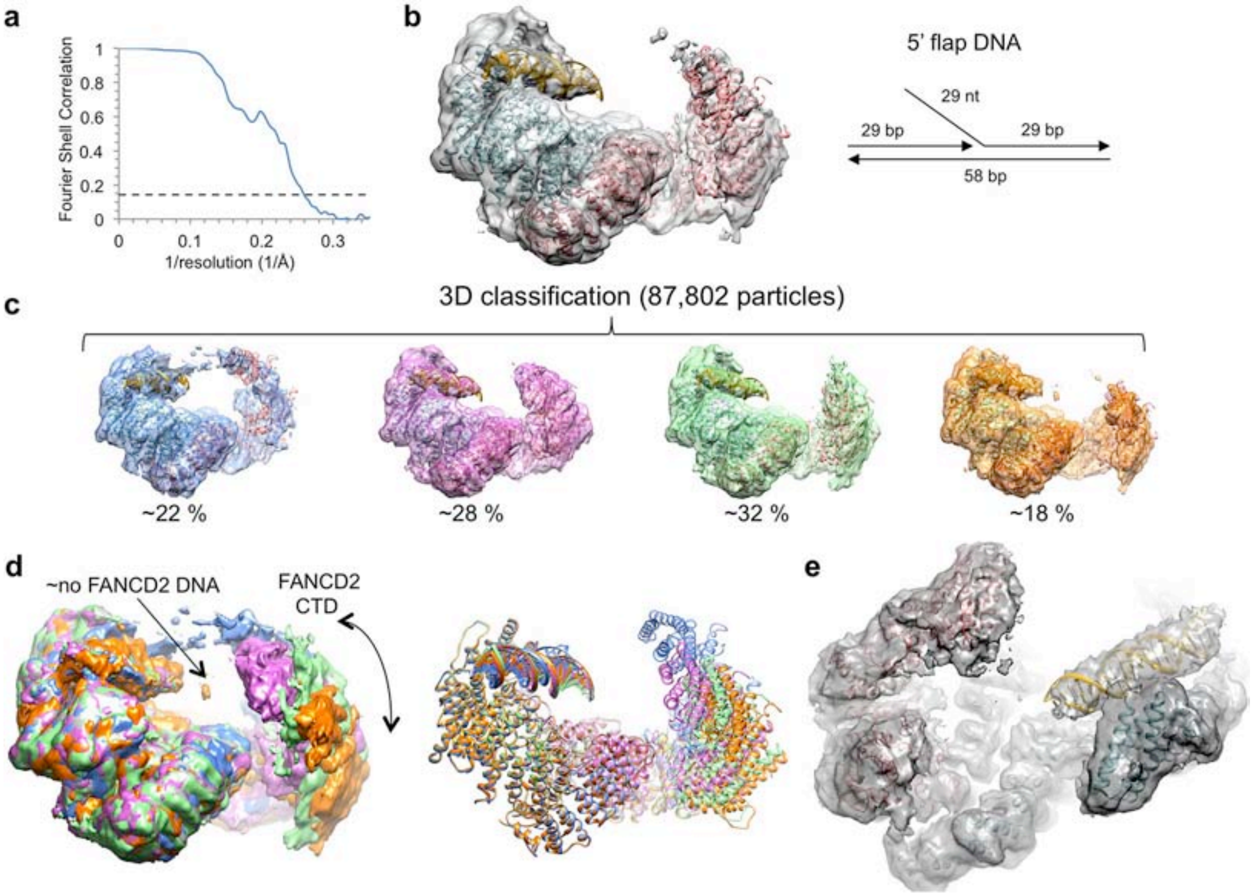


**Extended Data Figure 5 | Cryo-EM reconstruction of the non-ubiquitinated ID complex bound to 5' flap DNA.**

**a,** Graph shows gold-standard FSC plot between two independently refined half-maps for the consensus reconstruction with 87,802 particles. Dashed line marks the FSC cutoff of 0.143 that the FSC curve intersects at 4.0 Å.

**b,** Cryo-EM map from the consensus reconstruction prior to post processing. Also shown are cartoon representations of the FANCI (cyan), FANCD2 (pink) and FANCI-bound dsDNA (gold) from the ID-ICL DNA complex rigid-body fitted into the map. Schematic of the 5' flap DNA is shown to the right of the map.

**c,** 3D classification of the particles showing the conformational flexibility of FANCD2. Maps shown are after the particles from each 3D class were further refined in RELION to 4.7, 4.3, 4.3 and 4.9 Å, respectively. The maps are without temperature-factor sharpening. The ID complex and dsDNA, rigid body fitted into each map are also shown.

**d,** The five 3D classes are superimposed by aligning the FANCI-portion of each map (left), or of each pdb (right) colored as in **c**. The lack of FANCD2-bound dsDNA is indicated by the label "No FANCD2 DNA". The density of the FANCD2 CTD is significantly weaker and flatter than the similarly calculated maps of the of the ID-ICL DNA complex, suggesting increased mobility in the absence of appropriate DNA substrate. The curved arrow indicates the motion suggested by the 3D classification.

**e,** Close-up view of the best 3D class (pink in **d** after 3D refinement of the particles without post processing. Orientation is similar to Fig. 1c in the main text. Neither this map nor those of the other 3D classes have any evidence for a localized fork junction or for the 5' ssDNA flap, suggesting the 5' flap DNA binds to FANCI in multiple registers with no specificity for the junction. The maps for the Holliday junction (Extended Data Fig. 6) and fork (Extended Data Fig. 7) DNAs do show some bifurcation at one end of the duplex, but this is likely due to their shorter duplexes of 20 and 21 bp, respectively, being just long enough to fill the FANCI groove.

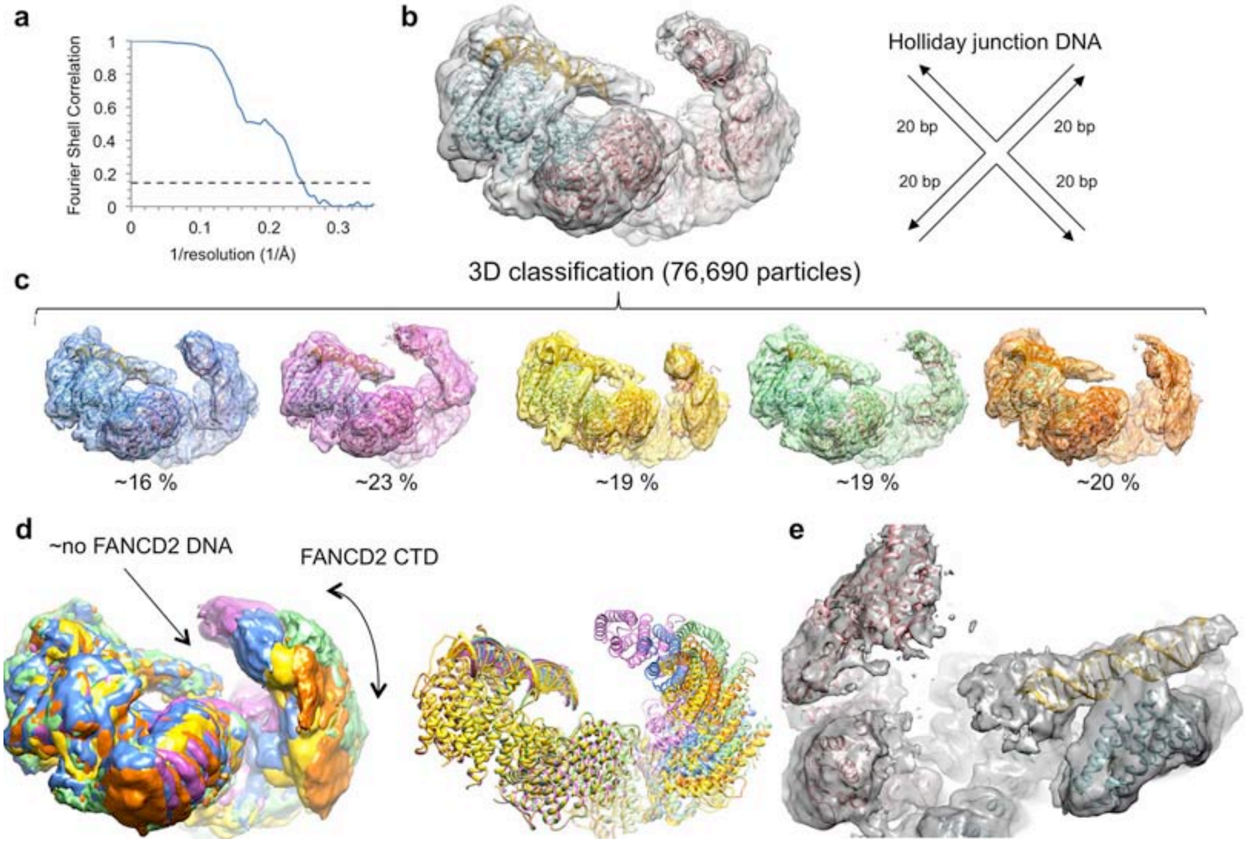


**Extended Data Figure 6 | Cryo-EM reconstruction of the non-ubiquitinated ID complex bound to Holliday junction DNA.**

**a,** Gold-standard FSC plot between two independently refined half-maps for the consensus reconstruction with 76,690 particles, with an FSC value of 0.143 (dashed line) at 4.1 Å.

**b,** Cryo-EM map from the consensus reconstruction prior to post processing. Also shown are cartoon representations of the FANCI (cyan), FANCD2 (pink) and FANCI-bound dsDNA (gold) from the ID-ICL DNA complex rigid-body fitted into the map. Schematic of the Holliday junction DNA used is shown to the right of the map.

**c,** 3D classification of the particles showing the conformational flexibility of FANCD2. Maps shown are after the particles from each 3D class were further refined in RELION to 6.9, 4.7, 6.6, 4.7 and 6.5 Å, respectively. The maps are without temperature-factor sharpening. The ID complex and dsDNA, rigid body fitted into each map are also shown.

**d,** The five 3D classes are superimposed by aligning the FANCI-portion of each map (left), or of each pdb (right) colored as in **c**.

**e,** Close-up view of the best 3D class (green in **d**) after 3D refinement of the particles prior to post processing. Orientation similar to Fig. 1c in the main text.

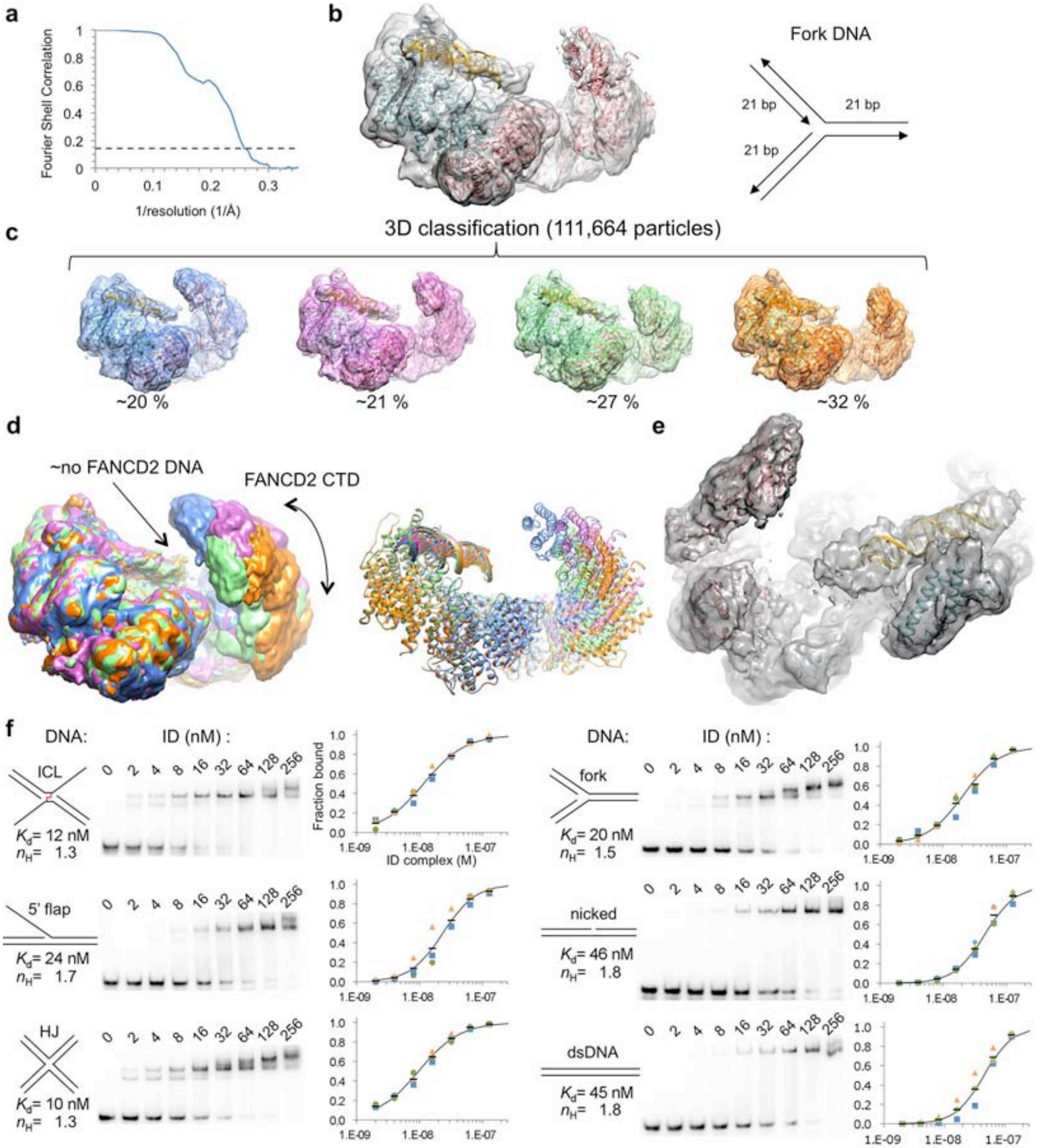
a
Fourier Shell Correlation
1/resolution (1/Å)
b
Fork DNA
21 bp
21 bp
21 bp
c
3D classification (111,664 particles)
~20 %
~21 %
~27 %
~32 %
d
~no FANCD2 DNA
FANCD2 CTD
e
f
DNA:
ID (nM) :
0 2 4 8 16 32 64 128 256
ICL
$K_d$= 12 nM
$n_H$= 1.3
Fraction bound
ID complex (M)
5' flap
$K_d$= 24 nM
$n_H$= 1.7
HJ
$K_d$= 10 nM
$n_H$= 1.3
fork
$K_d$= 20 nM
$n_H$= 1.5
nicked
$K_d$= 46 nM
$n_H$= 1.8
dsDNA
$K_d$= 45 nM
$n_H$= 1.8

**Extended Data Figure 7 | Cryo-EM reconstruction of the non-ubiquitinated ID complex bound to replication fork DNA and electrophoretic mobility shift assay data.**

**a,** Gold-standard FSC plot between two independently refined half-maps for the consensus reconstruction with 111,664 particles, with an FSC value of 0.143 (dashed line) at 3.9 Å.

**b,** Cryo-EM map from the consensus reconstruction prior to post processing. Also shown are cartoon representations of the FANCI (cyan), FANCD2 (pink) and FANCI-bound dsDNA (gold) from the ID-ICL DNA complex rigid-body fitted into the map. Schematic of the replication fork DNA used is shown to the right of the map.

**c,** 3D classification of the particles showing the conformational flexibility of FANCD2. Maps shown are after the particles from each 3D class were further refined in RELION to 4.8, 4.7, 4.6 and 4.4 Å, respectively. The maps are without temperature-factor sharpening. The ID complex and dsDNA, rigid body fitted into each map are also shown.

**d,** The five 3D classes are superimposed by aligning the FANCI-portion of each map (left), or of each pdb (right) colored as in **c**.

**e,** Close-up view of the best 3D class (orange in **d**) after 3D refinement of the particles prior to post processing. Orientation similar to Fig. 1c in the main text.

**f,** Electrophoretic mobility shift assay (EMSA) of the ID complex binding to the indicated $^{32}P$-labeled DNA substrates (0.5 nM) in the presence of 1.4 μM unlabeled, 20 bp dsDNA as nonspecific competitor. The plots with a logarithmic X-axis show fraction bound in three repetitions of each experiment (blue, green, orange markers), and their mean value (black dash). Each binding isotherm fits a Hill slope model significantly better than a non-competitive binding model, even after excluding the highest protein concentration reactions where multiple shifted bands are apparent, and also in the absence of nonspecific competitor DNA (not shown). It is likely this reflects, at least in part, the dissociation of the ID complex into monomeric FANCI and FANCD2 at the lower protein concentrations. A binding curve (black line) simulated with the indicated $K_d$ and Hill coefficient ($\eta_H$) values is shown on each plot.

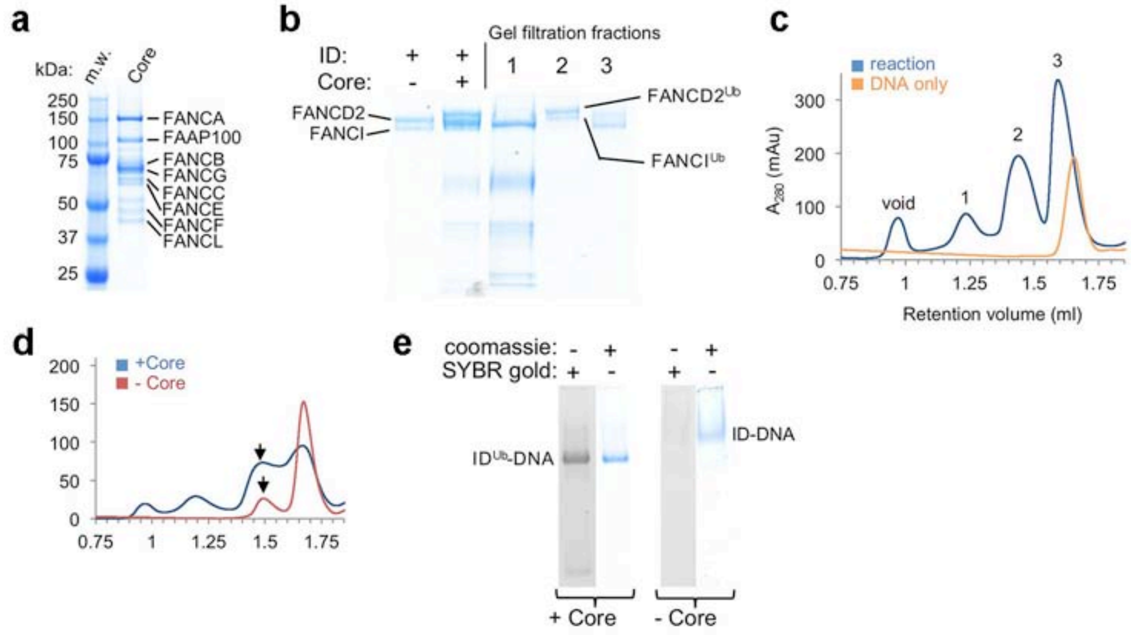


**Extended Data Figure 8 | Reconstitution of ID ubiquitination in vitro.**

**a,** SDS-PAGE gel of the purified FA core complex. M.w.: molecular weight markers with their mass labeled; Core: FA Core complex with the constituent proteins labeled.

**b,** SDS-PAGE of the ubiquitination reaction of the ID complex in the presence of a 58 bp nicked-DNA molecule and of three peaks from the fractionation of the reaction products on a Superose 6 gel-filtration column shown in b.

**c,** Gel-filtration chromatography of the ubiquitination reaction products (blue plot) and of the DNA-only control (orange plot). The fraction marked 1 contains the core complex, fraction 2 the complex of uniquitinated FANCI and ubiquitinated FANCD2, and fraction 3 contains monomeric non-ubiquitinated FANCI and FANCD2, as well as the overlapping peak of excess DNA.

**d,** Comparison of the gel-filtration chromatography profiles of the ID ubiquitination reaction (blue plot) and of non-ubiquitinated ID (red plot), both at 8 μM, in the presence of 16 μM ICL-DNA and 100 mM NaCl.

**e,** 4 % TBE gels of the gel-filtration fractions marked with an arrow in **d** stained for protein with Coomassie Blue and for DNA with SYBR Gold. The DNA band is faint in the non-ubiquitinated ID-DNA fraction due to its low concentration.

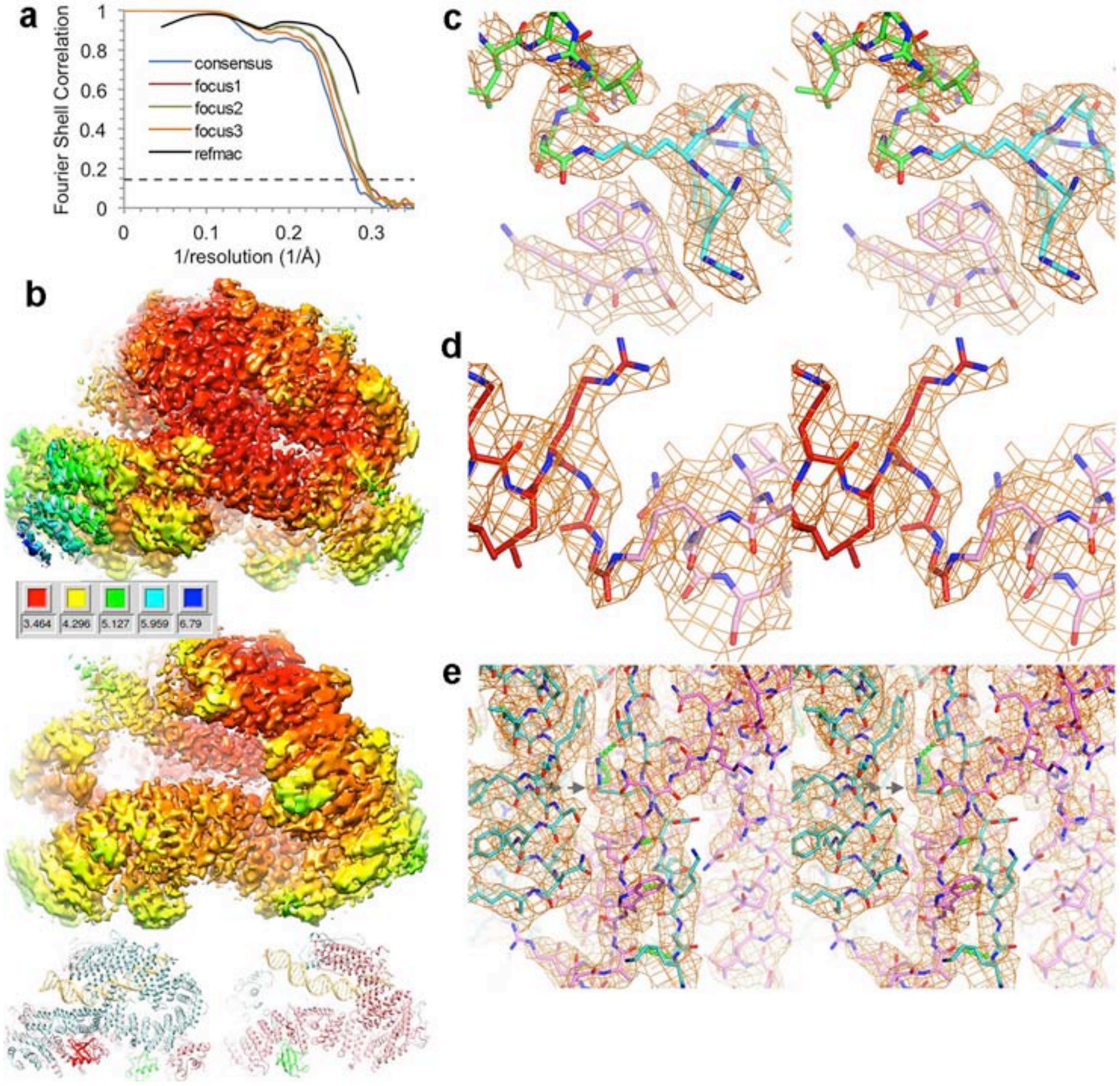


**Extended Data Figure 9 | Cryo-EM reconstruction of the ID$^{Ub}$ complex bound to nicked DNA.**

**a,** Graph shows gold-standard FSC plots between two independently refined half-maps for the consensus reconstruction of ID$^{ub}$ bound to nicked dsDNA (blue curve), and of the focused refinements with three different masks as described in Methods (focus 1, 2, and 3 in red, green and orange, respectively). The FSC curve for the final refined model versus the composite map combining the cryo-EM maps of the three focused reconstructions in REFMAC5 is shown in black. Dashed line marks the FSC value of 0.143.

**b,** Local resolution estimation, done with RELION3, of the ID$^{Ub}$-nicked DNA consensus reconstruction at 3.57 Å, viewed from the side of FANCI (top map, left pdb at bottom), or from the side of FANCD2 as in Fig. 2b (bottom map, right pdb). Maps are colored according to the resolution indicated in the inset (red 3.464, yellow 4.296, green 5.127, cyan 5.959, and blue 6.79 Å).

**c,** Stereo view of the 3.48 Å cryo-EM density of the isopeptide bond between the ubiquitin Gly76 C atom and the $N_{\zeta}$ atom of Lys523 of FANCI from the post-processed reconstruction of the focused refinement with mask 2. $Ub^{I}$ is green, FANCI cyan. O and N atoms are colored half-bonded red and blue, respectively, for both proteins. Also shown is FANCD2 Trp182 (pink) that packs with Lys523.

**d,** Stereo view of the 3.48 Å cryo-EM density of the isopeptide bond between the ubiquitin Gly76 C atom and the $N_{\zeta}$ atom of Lys561 of FANCD2 from the post-processed reconstruction. $Ub^{D2}$ is dark red, FANCD2 pink.

**e,** Stereo view of the 3.44 Å cryo-EM density of the zipper β sheet of the ID$^{Ub}$ complex. FANCI is cyan and FANCD2 pink. The arrow points to FANCI Arg1285 that is mutated to Gln in Fanconi Anemia. Select hydrogen bonds (made by the β sheet and by Arg1285) are shows as green dotted lines.

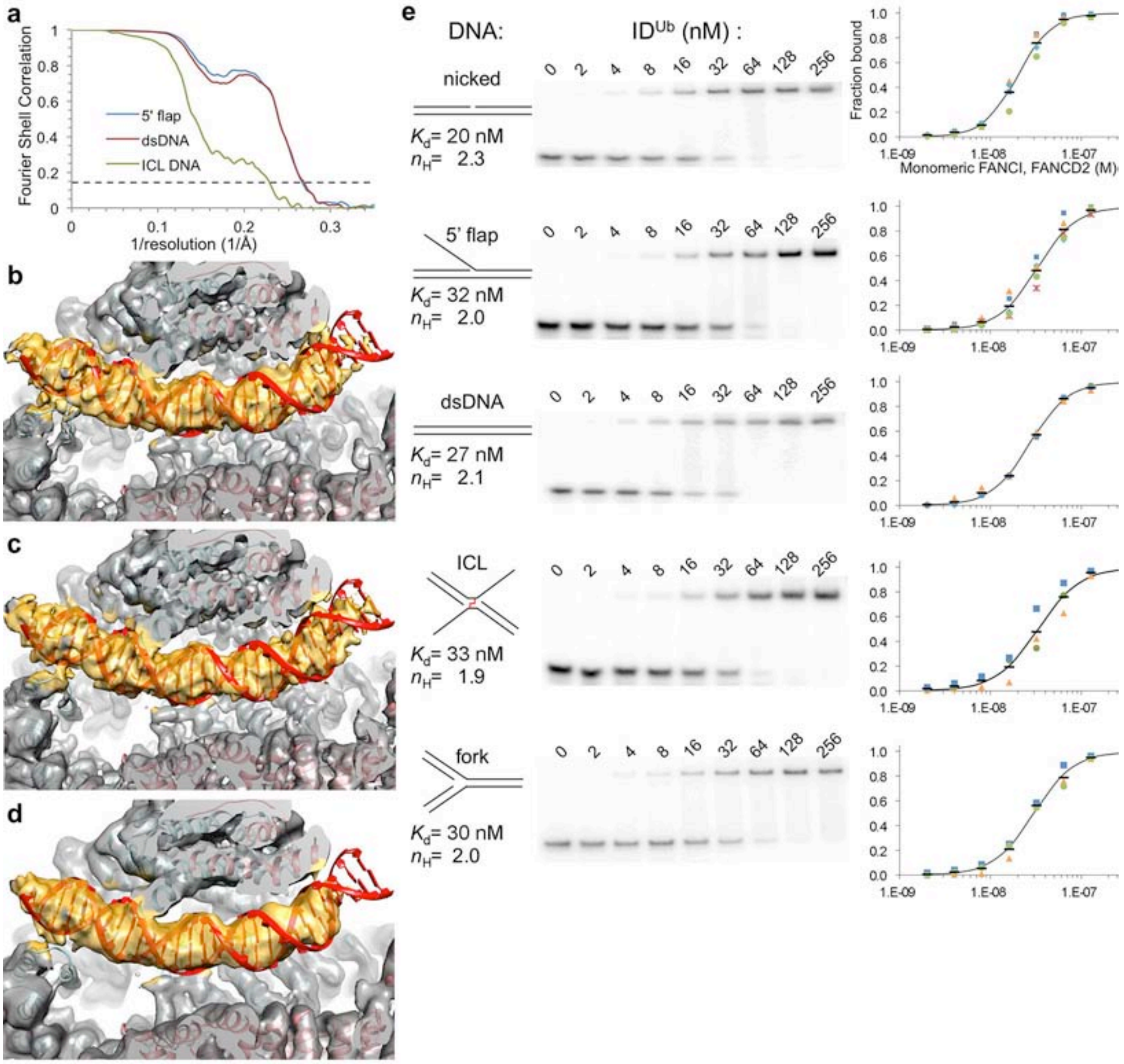


**Extended Data Figure 10 | Cryo-EM reconstructions of the ID$^{Ub}$ complex bound to 5' flap DNA, dsDNA, and ICL DNA and electrophoretic mobility shift assay data.**

**a,** Gold-standard FSC plots for the consensus reconstructions of ID$^{Ub}$ bound to the other DNA molecules discussed in text. The blue curve is the 3.8 Å reconstruction from 98,750 particles of ID$^{Ub}$ bound to 5' flap DNA (same DNA as the non-ubiquitinated complex of Extended Data Fig. 5b, the red curve is the 3.8 Å reconstruction from 85,078 particles of ID$^{Ub}$ bound to 58 bp dsDNA, and the green curve is the 4.4 Å reconstruction from 28,519 particles of $ID^{Ub}$ bound to the ICL-DNA of Extended Data Fig. 1b.

**b-d,** Cryo-EM reconstruction of $ID^{Ub}$ bound to **b** 5' flap DNA (two 29 bp duplexes flanking the flap) **c** dsDNA (58 bp) and **d** ICL DNA. Maps shown are prior to post-processing, at the resolutions indicated in **b**. Orientation similar to Fig. 3c, with the FANCD2 portion above the plane of the figure clipped. The model shown is the $ID^{Ub}$-nicked DNA structure that was fit as a single body into each map. We did not refine the coordinates for these reconstructions, yet the DNA density for each overlaps well with the nicked DNA model. The DNA density of the ICL DNA substrate is truncated at the FANCD2 end of the DNA, consistent with the canonical dsDNA arms of the 5' flap (29 bp) and ICL DNA (21 bp) substrates binding to $ID^{Ub}$.

**e,** EMSA of the equimolar mixture of the mono-ubiquitinated $FANCI^{Ub}$ and $FANCD2^{Ub}$, each at the indicated concentrations, binding to the $^{32}P$-labeled DNA substrates (0.5 nM) shown schematically. The plots with a logarithmic X-axis show fraction bound in three repetitions of each experiment, except for nicked and flap DNA, which were repeated 4 and 5 times, respectively, and their mean value (black dash). As with the non-ubiquitinated complex, the binding isotherms fit a Hill slope model best. We presume this reflects the hetero-dimerization of the monomeric $FANCI^{Ub}$ and $FANCD2^{Ub}$, as the data can be fit with similar residuals by including a dissociation constant for $FANCI^{Ub}$-$FANCD2^{Ub}$ hetero-dimerization in a non-competitive DNA-binding model (not shown). A binding curve (black line) simulated with the indicated $K_d$ and Hill coefficient ($h_H$) values is shown on each plot.

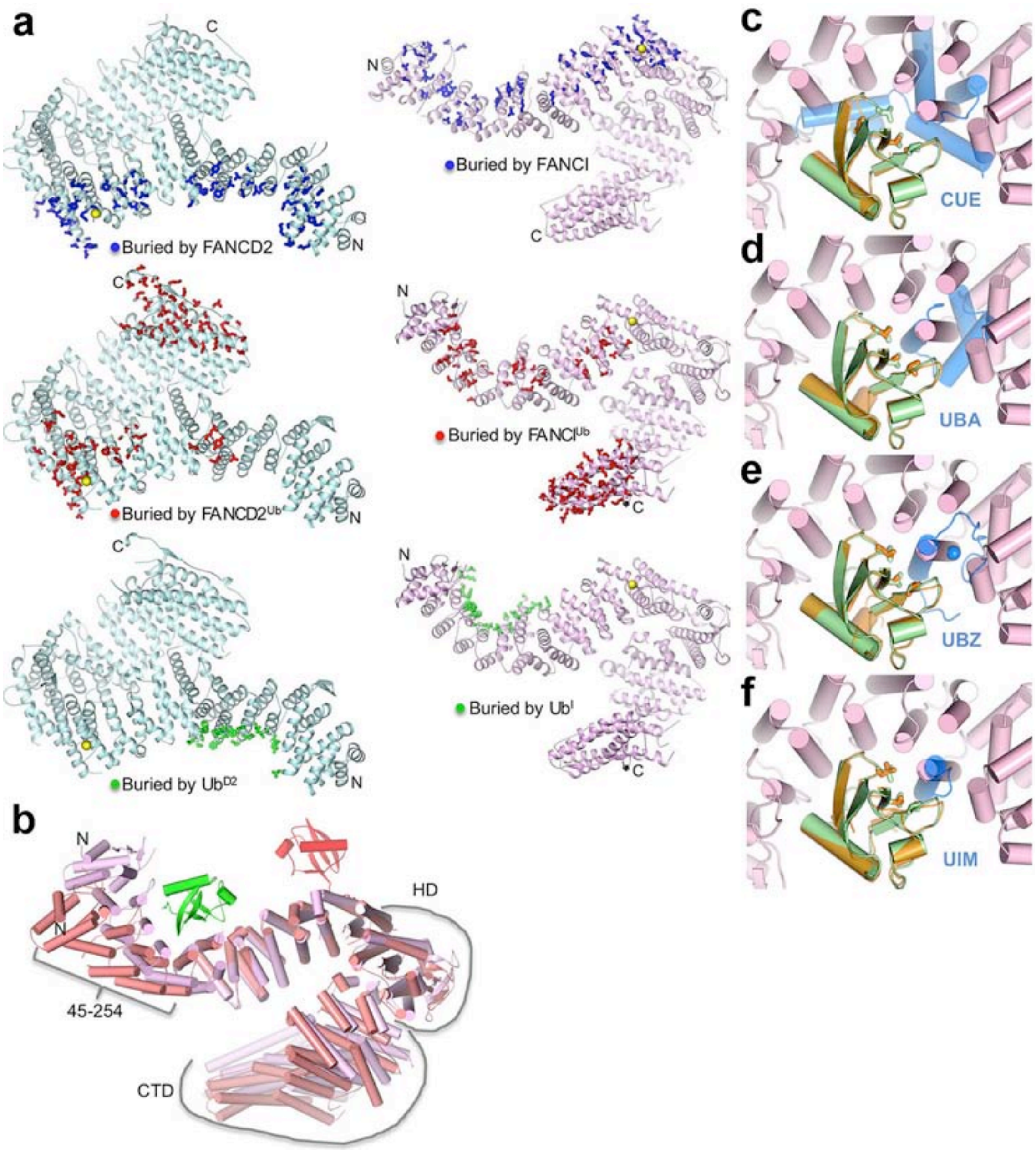


**Extended Data Figure 11 | Ubiquitination induces conformational changes and alternative FANCI-FANCD2 contacts.**

**a,** Cartoon representation of monomeric FANCI (left column, cyan) and FANCD2 (right column, pink) proteins showing residues (thick sticks) with a reduction in solvent accessibility due to interactions between FANCI and FANCD2 (top pair, blue sticks), between $FANCI^{Ub}$ and $FANCD2^{Ub}$ (middle pair, red sticks), and between each FANC protein and the ubiquitin on the other paralog (bottom pair, green sticks). The N and C termini are labeled for each. Yellow spheres indicate the ubiquitination site of each FANC protein.

**b,** FANCD2 from the ID complex (salmon) is superimposed on that of $ID^{Ub}$ (pink) by aligning the $2^{nd}$ half of their NTDs (residues 255-587; 1.8 Å r.m.s.d. for 308 Cα atoms), a segment that changes little on ubiquitination. $Ub^{D2}$ that is covalently attached to FANCD2 is in red, and the $Ub^{I}$ with which it packs in green. The N-terminal portion of the NTD, which rotates by 38° towards FANCI as a rigid body (residues 45 to 254, 0.68 Å r.m.s.d. for 202 $C_a$) is approximately marked by a bracket. This rotation at the center of the $Ub^{I}$ binding site allows FANCD2 to better embrace $Ub^{I}$, and also to interact with FANCI (NTD-HD junction, residues 529 to 593), the latter involving similar residues on FANCD2 but mostly different ones on FANCI. Similarly marked is the HD domain (residues 604-928) that rotates relative to the NTD by 15°. Additional tilting of helices within the HD domain results in the CTD (residues 929 to C-termini also marked) that follows being rotated by 20° degrees relative to the invariant portion of the NTD.

**c-f,** The FANCI and FANCD2 ubiquitin binding structural elements are distinct from commonly occurring ubiquitin-binding domains. Superposition of the $Ub^{I}$ (green) bound to $FANCD2^{Ub}$ (pink) on the ubiquitin (orange) bound to: **c,** the dimeric Vps29 CUE domain (blue) from PDBID 1P3Q, **d,** the Cbl-b UBA domain (PDBID 2OOP), **e,** the UBZ domain of Faap20 (PDBID 3WWQ), and **f,** the UIM domain of Vps27 (PDBID 1Q0W). The ubiquitin hydrophobic patch residues (Leu8, Ile44, and Val70) are shown in stick representation for both ubiquitin molecules in each figure. The blue sphere in **e** is the Zn atom of the UBZ domain. Orientation similar to that of Fig. 4a. It has been suggested that FANCD2 shares sequence homology with the CUE domain[31]. While the 47-residue region of proposed homology (residues 191-237) partially overlaps the Ub-binding site, its structure is unrelated to the CUE domain, and Ub binding by FANCD2 is distinct.

**Extended Data Table 1. Cryo-EM data collection, model refinement and validation statistics.**

| | ID-ICL DNA | ID$^{Ub}$-nicked DNA |
|---|---|---|
| **Data collection and processing** | | |
| Voltage (kV) | 300 | 300 |
| Electron exposure (e–/Å$^2$) | 65.6 | 65.6 |
| Pixel size (Å) | 1.089 | 1.089 |
| Particle images (no.) | 231,943 | 301,058 |
| Defocus range (μm) | 1.0-3.2 | 1.0-3.2 |
| Map resolution (Å)$^{\#}$ | 3.40 | 3.57 |
| **Refinement** | Focused maps | Multi-body maps |
| Resolution (Å) | 173.31 - 3.35 | 167.62 - 3.48 |
| Average FSC | 0.835 (0.591)$^*$ | 0.864 (0.619)$^*$ |
| $R_{work}$ (%) | 34.25 | 31.32 |
| Non-hydrogen atoms | | |
| Protein | 18,517 | 20,026 |
| DNA | 1,538 | 1,189 |
| B factors (Å$^2$) | | |
| Protein | 169.4 | 154.3 |
| DNA | 292.3 | 230.5 |
| R.m.s. deviations | | |
| Bond lengths (Å) | 0.011 | 0.011 |
| Bond angles (°) | 1.42 | 1.59 |
| B factors main chain (Å$^2$) | 2.9 | 2.6 |
| B factors side chain (Å$^2$) | 0.8 | 3.6 |
| Ramachandran plot | | |
| Disallowed (%) | 0.04 | 0.16 |
| Favored (%) | 93.97 | 94.69 |
| Validation | | |
| MolProbity score | 1.62 | 1.59 |
| Clashscore | 4.35 | 3.28 |
| Rotamer outliers (%) | 0.75 | 1.39 |

$^{\#}$Resolution at FSC=0.143 is for the consensus reconstruction for each data set.

$^*$Values in parenthesis are for the highest resolution shell.

**Extended Video 1.**

Movie illustrates the top mode of conformational flexibility of the FANCD2 CTD in the non-ubiquitinated ID-ICL DNA complex. The ten maps of the movie were generated by the relion_flex_analyze (RELION-3) program for the top component from the PCA of the multi-body refinement angles described in the Extended Data Fig. 3a legend. The model from the consensus reconstruction is shown in cartoon representation and colored as in Fig. 1a.

**Extended Video 2.**

Movie illustrates a second mode of conformational flexibility in the FANCD2 CTD of the non-ubiquitinated ID-ICL DNA complex. The ten maps are for the second PCA component (Extended Data Fig. 3b).

30 Joo, W. *et al.* Structure of the FANCI-FANCD2 complex: insights into the Fanconi anemia DNA repair pathway. *Science* **333**, 312-316 (2011).

31 Rego, M. A., Kolling, F. W. t., Vuono, E. A., Mauro, M. & Howlett, N. G. Regulation of the Fanconi anemia pathway by a CUE ubiquitin-binding domain in the FANCD2 protein. *Blood* **120**, 2109-2117 (2012).
